# Multiscale light-sheet organoid imaging framework

**DOI:** 10.1101/2021.05.12.443427

**Authors:** Gustavo de Medeiros, Raphael Ortiz, Petr Strnad, Andrea Boni, Franziska Moos, Nicole Repina, Ludivine Chalet Meylan, Francisca Maurer, Prisca Liberali

**Author notes:** Correspondence should be addressed to P.L.

## Abstract

Organoids provide an accessible *in-vitro* system to mimic the dynamics of tissue regeneration and development. However, long-term live-imaging of organoids remains challenging. Here we present an experimental and image-processing framework capable of turning long-term light-sheet imaging of intestinal organoids into digital organoids. The framework combines specific imaging optimization combined with data processing via deep learning techniques to segment single organoids, their lumen, cells and nuclei in 3D over long periods of time. By linking lineage trees with corresponding 3D segmentation meshes for each organoid, the extracted information is visualized using a web-based “Digital Organoid Viewer” tool allowing unique understanding of the multivariate and multiscale data. We also show backtracking of cells of interest, providing detailed information about their history within entire organoid contexts. Furthermore, we show cytokinesis failure of regenerative cells and that these cells never reside in the intestinal crypt, hinting at a tissue scale control on cellular fidelity.

## Introduction

During adult life, organs such as the intestine are challenged to diverse environmental conditions, requiring the tissue to be robust and yet plastic. For instance, during regeneration after damage, surviving cells need to carefully orchestrate a fast and robust regrowth process in coordination with proper shape recovery as well as functional and morphological remodeling. To best overcome the difficulties surrounding the study of these tissue dynamics in inner organs *in vivo*, organoids have become a powerful experimental method owing to their exceptional accessibility and manipulability^1-3^. For the case of the intestinal tract, intestinal organoids grown from single cells functionally recapitulate both the regenerative response of the intestinal epithelium as well as the homeostasis of the *in vivo* intestine^4^. Morphologically, they also recapitulate the main dynamics of crypt formation, making them a unique *in vitro* system^5,6,7, 8^.

Although with high degree of accessibility, performing live imaging of organoid growth remains a challenge, as it typically requires not only microscopy techniques capable of stable long-term imaging of several samples simultaneously, but also dedicated analysis and processing pipelines that can cope with complex imaging data. On the more technical imaging side, high-resolution multi-view light-sheet imaging has been used to track single cells in different embryo development settings ^9,10^, at the cost of low throughput imaging (usually 1-5 samples per imaging experiment). This is detrimental since the efficiency of organoid formation from single cells is particularly low (around 15% in the case of murine intestinal organoids^6^). Furthermore, previous work on the live recording of organoid dynamics has focused either on specific biological questions^6,8,11^, organoid wide phenotype-driven screening approaches^12-14^, or on specific isolated tools^15^ without a more generalised yet in-depth approach on light-sheet imaging and data analysis. In another work which aimed at creating a light-sheet organoid imaging platform^16^ the focus was mainly on the determination of culture-wide heterogeneities through a combination of both light-sheet and wide-field techniques. Although showing organoid diversity within the same culture, in-depth cellular multi-scale analysis for each organoid remained lacking.

To bridge the gap and provide quantitative information on organoid growth dynamics with in-depth cellular analysis, we here provide a unified multiscale light-sheet imaging framework tailored to live organoid imaging. Our framework incorporates optimized imaging, pre-processing, semi-automated lineage tracking, segmentation and multivariate feature extraction pipelines which provide multiscale measurements from organoid to single cell levels. By focusing on the development of intestinal organoids, we show that this holistic set of tools allows the combination of whole organoid and single cell features to be analysed simultaneously, having both lineage tree as well as spatial segmentation information presented in a clear and unified way. We demonstrate that our pipeline is compatible with fixation and immunolabeling after live imaging by tracking back cells positive for specific markers and compare the history of these cells with all other cells in the organoid. To facilitate the usage of the analysis and visualization tools, we have combined them into a unified set of tools we call LSTree^17^, built on a Luigi workflow and dedicated notebooks, allowing the different steps to run in a modular way. Further, we use our framework to dissect previously unknown biological insights on the role of polyploidy during intestinal organoid growth, and we propose a tissue level check point for tissue integrity that could start explaining the interplay between regeneration and cancer.

## Results

### Imaging framework for light-sheet microscopy of organoid growth

We developed a multiscale imaging framework that comprehends acquisition, pre-processing, automated tracking, segmentation with further feature extraction as well as visualization dedicated for 3D live imaging (**Fig. 1a**). In this work, we applied our framework to live intestinal organoid light-sheet recordings performed with a dual-illumination inverted light-sheet^6^ microscope, which utilizes a multi-positioning sample holder system (**Fig. 1b**). In order to image organoid development, we followed previously published protocols^6^, FACS sorting single cells (**Supplementary Figure 1a**) from mature organoids and mounting them as 5 uL mix drops with Matrigel on top of a ca. 50 um thick fluorinated ethylene propylene (FEP) foil, which are then covered in medium (**Fig 1c** left and **Methods Section**). To stabilize the imaging, we patterned the FEP foil used for mounting in order to create small wells (**Supplementary Figure 1b-d, Supplementary Note 1**), allowing better control of the sample position within the holder, while improving reproducibility of experiments by preventing drops from being washed away during medium change of fixation procedures. As previously demonstrated, the microscope we utilized is capable of imaging live intestinal organoids for long periods of time^6,8^, as well as acquiring time-lapses of mouse embryonic and gastruloid development^18,19^. However, one important drawback of the system was that the alignment of the illumination beams is done only once, prior to the experiment and irrespective of the position of the sample in the dish or holder. Although sufficient in certain situations (e.g. mouse embryo imaging), imaging of samples embedded and distributed inside a gel suffer from refractive index mismatch between water and Matrigel, as well as from the presence of other obstacles in the light-path (other organoids or debris) and from the curved shape of the sample holder itself. Therefore, to improve recording conditions in every individual sample, we developed a position dependent illumination alignment step. This allows to fine tune the alignment of each of the illumination sheets in respect to the detection plane for every sample position so that best image quality possible can be achieved throughout (**Fig 1b** right and **Supplementary Note 1**).

**Figure 1:**
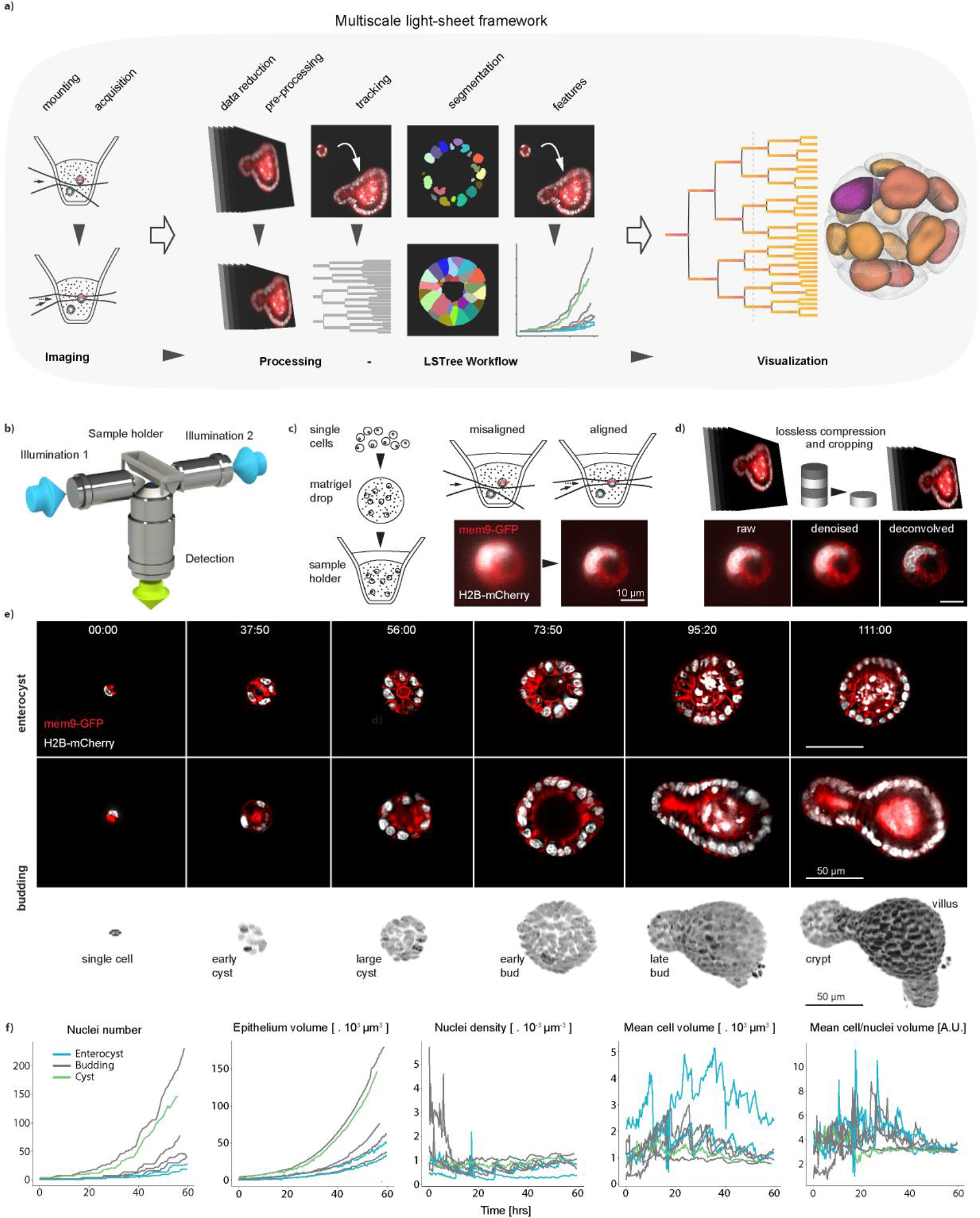
Acquisition of high-resolution 3D organoid images. **a)** Multiscale light-sheet imaging framework, depicting imaging stages, analysis workflow and visualization tool. **b)** Dual illumination inverted detection light-sheet objective configuration used in all of the recordings. **c)** Left: sample preparation is performed by mixing single cells dissociated from mature organoids with matrigel and depositing 5 uL drops on the light-sheet sample holder. Right: sample position dependent illumination alignment corrects for possible misalignments of the illumination beam in reaching organoids distributed inside the Matrigel drop, improving SNR. **d)** Upper row: minimization of storage needs done with on-the-fly compression and further cropping steps. Lower row: denoising and deconvolution steps further improve image quality. **e)** Time-lapse imaging of organoid growth. Top row shows still images of a growing enterocyst, whereas both lower rows show a stereotypical growth of a budding organoid as a cross sections as well as a projection in Paraview. **f)** Temporal evolution of nuclei number, epithelium volume, mean cell volume and the ratio between mean cell and mean nuclei volumes for all main 7 datasets considered in this work. Source data are provided as a Source Data file.

To minimize storage needs and improve SNR, acquired images are cropped using a dedicated tool that automatically corrects for 3D sample drifting (**Fig 1d** upper row). The cropped images may also be further pre-processed through denoising and deconvolution steps. Denoising is performed using the Noise2Void scheme^20^, with its output sent to a tensor-flow based image deconvolution^21^ (**Fig 1d** lower row) using measured PSFs from beads (**Supplementary Note 2** and **Methods Section**).

With these first modules at hand, we imaged organoids expressing Histone 2B and mem9 membrane peptide tagged with mCherry and GFP respectively, recording the growth and development of several organoids starting from single cells or 4-cell spheres (**Fig 1e** and **Supplementary Movie 1**) every 10 minutes throughout the course of around 4 days. The collected data comprised of organoids that form both budding and enterocyst phenotypes: whereas budding organoids grow from single cells into mature organoids with both crypt and villus structures, enterocysts, comprised of terminally differentiated enterocytes, do not have crypts as they do not develop Paneth cells required for the establishment of the stem cell niche, a necessary step for crypt formation^6,8^.

For the analysis, we initially performed single-cell partial semi-automatic tracking using the Fiji plugin Mastodon (https://github.com/mastodon-sc/mastodon) on 7 datasets. After that, we extracted features based on organoid and single cell segmentation and plotted this data over time (**Fig 1f** and **Supplementary Table 1**). For example, we noticed large variability in cell division synchronicity, as in some datasets the nuclei number growth over time loses the typical staircase-like behavior already early during the first day of recording. Although epithelium volume growth curves follow that of nuclei number, with the characteristic exponential behavior, nuclei density slightly increases over time. Mean cell volume showed characteristic mitotic peaks, with overall cell volume decrease over time, matching the increase in nuclei density. Interestingly, although initially cell to nuclei volume ratio vary, all datasets converge to common steady state values where the cell volume is ca. 3 to 4 times larger than the nuclear volume. We also observed a consistent change in organoid volume due to medium change during the live recordings (**Supplementary Figure 2**). As this initial assessment of our imaging data showed consistent and reproducible results, and to handle larger dataset more rapidly and consistently, we developed an integrated and automated approach to turn the imaging into digital organoids with a visualization tool.

### Dedicated image processing workflow

To make the entire analysis and visualization tools directly accessible, we incorporated all image processing and data analysis modules into a unified workflow named LSTree, having most processing and training steps implemented using Luigi based tasks, and the rest as jupyter notebooks for cropping and segmentation evaluation. The workflow along with juypter notebook and two example datasets are provided as a documented Github repository with step-by-step guide (see **Methods Section** and **Supplementary Notes 2-4** for more information).

In the first pre-processing step, the user selects which organoid needs to be cropped. This automatically generates minimal bounding boxes per time-point as well as global bounding box (**Fig 2a**). The workflow also has an interactive tool to review the crops and perform few manual corrections, e.g. to account for large displacement between consecutive frames (**Supplementary Figure 3**). Next, if needed, denoising and deconvolution of cropped and registered movies is performed as one combined step. Important to note that we chose to denoise and deconvolved our datasets as the image quality usually decays quite heavily at later timepoints. However, this is not a requirement, and the prediction models can also be trained based on good quality unprocessed datasets. More details on how to bypass the denoising and deconvolution steps are discussed through the example datasets provided in the GitHub documentation.

**Figure 2:**
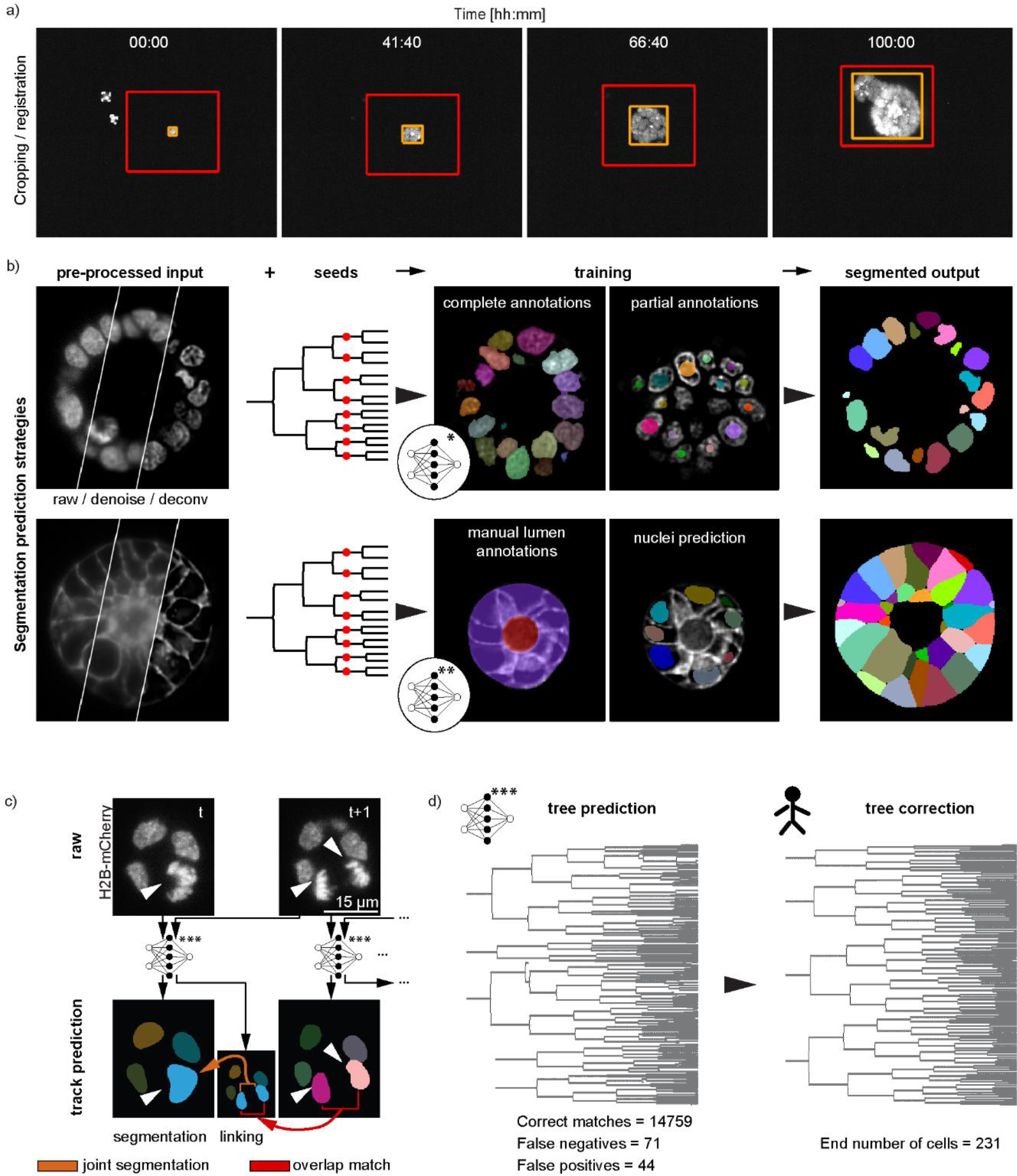
Cropping, segmentation and tree-prediction strategies underlying LSTree. **a)** Cropping of datasets is done in a semi-automatic way: selected object of interest is fitted with an orange (best fit for each particular timepoint) and a global red bounding boxes, which can be corrected in 3D. **b)** Nuclei and cell segmentation strategies. Upper row: denoised and deconvolved input data together with seeds from respective tracking are used as input into the network to predict nuclei volume. The network is trained with both complete and partial annotations. Lower row: Cell volume prediction follows similar input as for nuclei. Main difference is that this second network is trained with supervision of complete manual annotations of lumen and organoid along with the previously done nuclei predictions themselves. **c)** Strategy for prediction of lineage trees. Track predictions are done with each consecutive pair of frames. Each pair of frames enter the neural network and produce both the timepoint in question and a linking frame which is used to connect to the next timepoint via overlap match. Linking itself is done via joint segmentation. **d)** Example of predicted and corrected tree from a budding organoid dataset with the recording starting from two cells (recording 006).

For the segmentation of organoids, as well as their cells and nuclei, we adopted different segmentation strategies all relying on existing convolutional neural networks (**Fig 2b**). Our main initial motivation was to test whether we could incorporate the spatial information from the lineage trees spots for training segmentation models. To that end, we decided to use the RDCNet instance segmentation network as a base^22^, taking advantage of its inherent recursive architecture. First, nuclei are segmented in 3D following a deep learning model trained with a mix of complete and partial annotations. A small subset of the frames is fully annotated by manually expanding the labels to the full nuclei, whereas partial annotations rely on the initial tracking performed with Mastodon by drawing spheres at the position of tracked nuclei. (**Fig 2b** upper row). Jupyter notebooks for interactive visualization and correction of the predicted segmentation are also part of the framework and added onto the GitHub, which allows improving the model accuracy with minimal annotating time. To check whether this approach was valid, we compared the trained network output with randomly selected hand-annotated image volumes, yielding very good results (see **Supplementary Note 5** and **Supplementary Figure 4a-b**).

Motivated by the initial results with nuclei segmentation based on sparse annotations, we took a similar approach for cell segmentation. To this end, organoid and cell segmentation also use RDCNet and leverages the pre-computed nuclei segmentation to avoid manual annotations of individual cells. At the same time, we added a constraint based on lumen and epithelium segmentation, to avoid that cell labels spread outside of the epithelial layer. To subdivide the epithelium mask into cells, the previously segmented nuclei are used as partial cell annotations under the assumption that they are randomly distributed within the cell compartment (**Fig 2b** lower row, **Supplementary Note 3**). Finally, in addition to the segmentation volumes and nuclei number (**Fig. 1e**), several different features are extracted such as nuclear distance to apical/basal membranes, fluorescence intensity, distance to parent node and number of neighbors per cell (For a complete list of features with short explanations see **Supplementary Table 1**).

### Deep learning model for automated lineage tracing

Although suitable for estimating lineage trees for few datasets, semi-automated tracking of many datasets with the Mastodon Fiji plugin can be time consuming, as different datasets may require different setting parameters often break when cells are too packed or with low signal-to-noise. To significantly improve this process, we trained and refined a deep learning model on the available tracked datasets aiming at automatic generation of candidate trees that only require minimal corrections (**Fig 2c,d, Supplementary Note 3**). To avoid usage of tracing algorithms that enforce a complex set of rules^23-25^, we developed a joint segmentation-tracking approach that simultaneously predicts matching nuclei labels on 2 consecutive frames. To this end, we extended the RDCNet instance segmentation model to predict pseudo 4D labels (3D convolutional network with time axis as an additional image channels) mapping correspondences between nuclei in 2 consecutive frames (**Fig 2c**). Predicting linked nuclei segmentations has the advantage to enforce constancy over the entire nuclear volume rather than relying on an ambiguous center, as well as implicitly enforcing rules such as minimum cell distance or plausible nuclei volume constraints in a data-driven manner. This method keeps the number of manual hyper-parameters tuning to a minimum and can be improved over time as more validated and corrected datasets are incorporated in the training set. In a complementary manner, this method can be used together with other deep learning strategies such as Elephant in a complementary and modular manner, in which curated trees via Elephant can be used for training of more generalized tracking models based on RDCNet, or even directly used for nuclei/cell segmentation training/prediction.

To assemble the predicted tree in the framework, nuclei labels in each frame are connected to their parents in the previous frame by finding the linking label with the maximum overlap (**Fig 2c, Supplementary Figure 5**). The predicted tree is then saved with the structure of a MaMuT.xml track file, which can be then imported into Mastodon for further correction if necessary (**Fig 2d**). As a direct consequence of the joint segmentation, additional information, such as the nuclei volume, can be overlaid on the predicted trees to aid in the curation process (**Supplementary Figure 5**, Supplementary **Note 3**). For instance, jumps in nuclear volume highlight positions where tracks should be merged or split. The manual curation time ranges from minutes to a couple hours on the most challenging datasets (e.g. low SNR images, abnormal nuclei shape). In summary, the here developed lineage tree prediction approach allows high quality prediction of intestinal organoid lineage trees with long tracks spanning multiple division cycles (up to 5 generations in this work) enabling tracked data to cross spatiotemporal scales. To further challenge our tracking prediction strategy, we have also tested it outside of our main focus on live imaging of intestinal organoid, and used trained models to validate prediction accuracy on mouse embryo datasets from published work (**Supplementary Figure 4c-e**, and discussions on **Supplementary Note 5**), also comparing it to output from trained Elephant Tracker models (all trained models can be found in the **Supplementary Software**).

### Digital organoid viewer

With the lineage trees and the deep learning 4D segmentation of organoid, lumen, cells and nuclei at hand, we developed a multiscale digital organoid viewer to explore and perform in-depth data mining. The viewer combines both lineage trees and segmented meshes, facilitating the direct comparison of different features within a multiscale digital organoid framework. We have added it to our LSTree Github repository along with example data, also including the possibility to overlay recorded images with the corresponding meshes allowing a direct inspection of the predicted segmentation (**Fig 3a, Supplementary Movie 2**). As can be seen through the example datasets present in the repository, this interactive viewer allows associated features to be displayed, selected nodes to be interactively highlighted on the meshes, and color coding of both trees and meshes to be be assigned independently. This way same or complementary features can be visualized at once (All currently extracted features are discussed in **Supplementary Note 4** and **Supplementary Table 2**)

**Figure 3:**
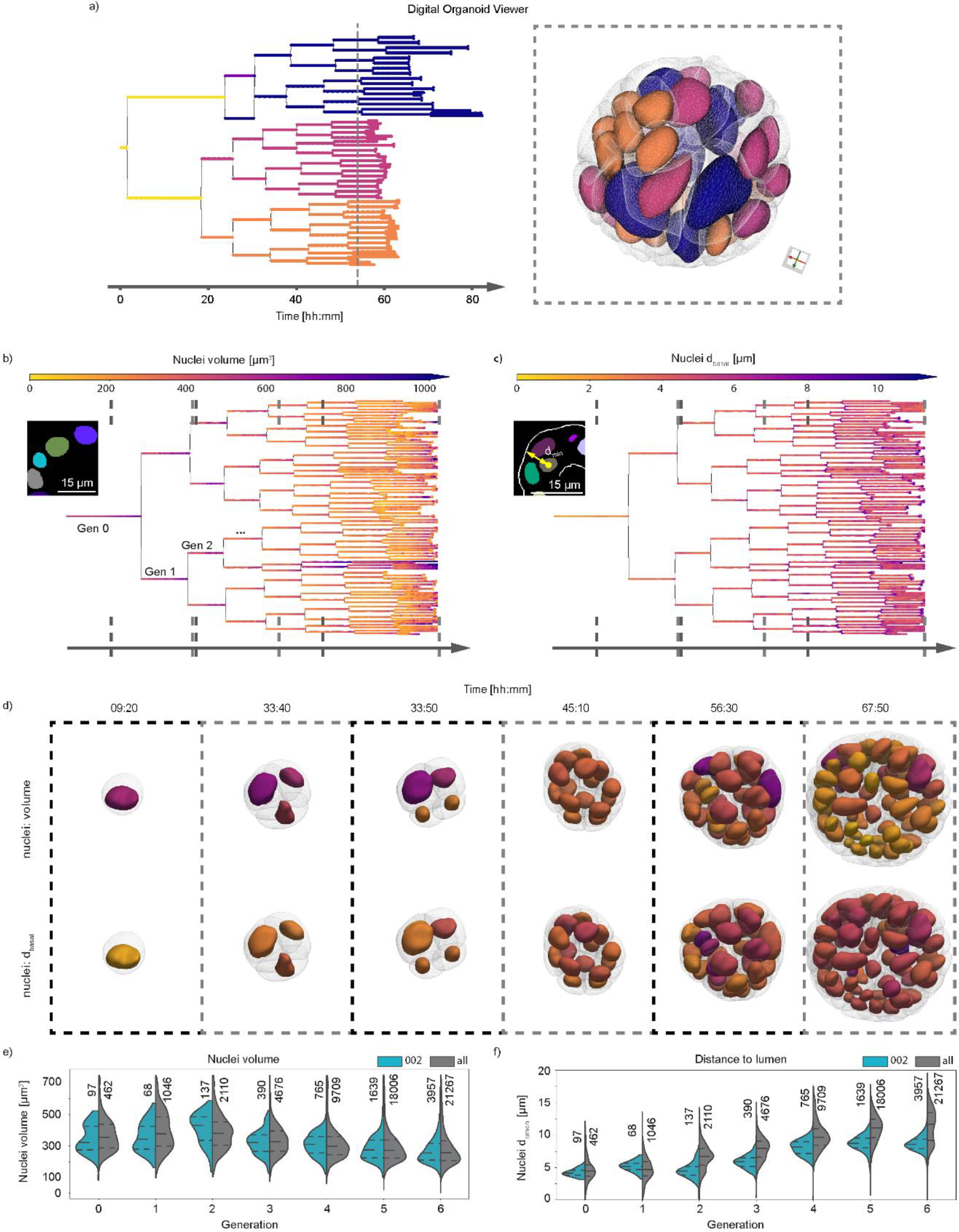
Digital Organoid Viewer. **a)** Digital organoid viewer is a web-based tool that shows both lineage trees (left) and respective segmented nuclei and cell meshes (right) simultaneously. Color coding of of each data representation can be done in a combined or complementary manner. Here depicted is a color coding based on generation 2, with each of the four cells acquiring different colors being propagated further in time. **b)** Overlay of the nuclei segmentation as predicted in onto the lineage tree. **c)** Overlay of distance to basal membrane onto the lineage tree. **d)** Visualization of the calculated meshes from nuclei and cell segmentations, overlaying the corresponding values (with corresponding color map) of the features presented in b) and c). The time points chosen are shown via dashed lines on each tree in b) and c). **e**,**f)** Extracted values for nuclei volume **(e)** and for distance to apical membrane **(f)** of the here exemplified dataset (pull, recording 002) against all datasets (all 7 recordings) analyzed. The dashed lines inside each plot correspond to the first and third quartile of the values from all of the datasets, with the median as the dashed line in between them. Each generation spans over the full cell cycle of all nuclei considered, and all nuclei corresponding to each plot are shown slightly above each violinplot. Source data are provided as a Source Data file.

As an example of the image-analysis and visualization tools presented in the framework, nuclear volume quantifications can be evaluated directly onto the tree of a specific dataset (**Fig 3b**). Using this approach, it is possible to observe and quantify how much nuclei volumes change with each generation and over time, with the smallest volumes observed right after division. Similarly, we observe that the nuclear distance to the basal membrane (**Fig 3c**) increases due to interkinetic nuclear migration towards the apical side. Combining the same visualization procedure with the segmented meshes, we render the nuclei or cells in 3D, using the same color-coding as for feature on the trees (**Fig 3d**). Last but not least, we also compare the extracted features against the general trend from all other datasets combined, allowing us a direct evaluation of variability across experiments (**Fig 3e,f**), evaluating the increased distancing of nuclei from the apical membrane, a known effect due to epithelial polarization (**Fig 3f**).

In summary, this is a unique set of tools embedded under the same workflow which allows not only multiscale segmentation of organoids along with lineage tree predictions, but also the simultaneous visualization of both trees and segmented meshes into a unified web-viewer. All steps of the process are implemented to keep storage, memory, and manual tuning requirements to a minimum, making this a powerful and yet easily accessible part of the light-sheet framework.

### Functional imaging through fixation and backtracking

Next, we analyzed functional information on the tracked cells and organoids contained in the lineage trees. Although our imaging framework allows the visualization and quantification of a large number of features at the cellular and organoid levels throughout organoid growth, functional information remains dependent on fluorescent reporter organoid lines. The easiest way to theoretically approach this is to perform stable multicolor live imaging for long periods of time. However, overlapping emission/excitation spectra limits the total number of fluorescent reporters and concerns regarding interference with the normal cell function, signal-to-noise sensitivity for low abundant proteins, photostability and general phototoxicity due to laser illumination limit the use of fluorescent reporters. To overcome this, we fixed and added an immunolabelling steps at the end of the live recordings to assess the end state of the cells. Then we tracked the immunolabelled cells back through the lineage tree (**Fig 4a**), using LSTree for further visualization and analysis.

**Figure 4:**
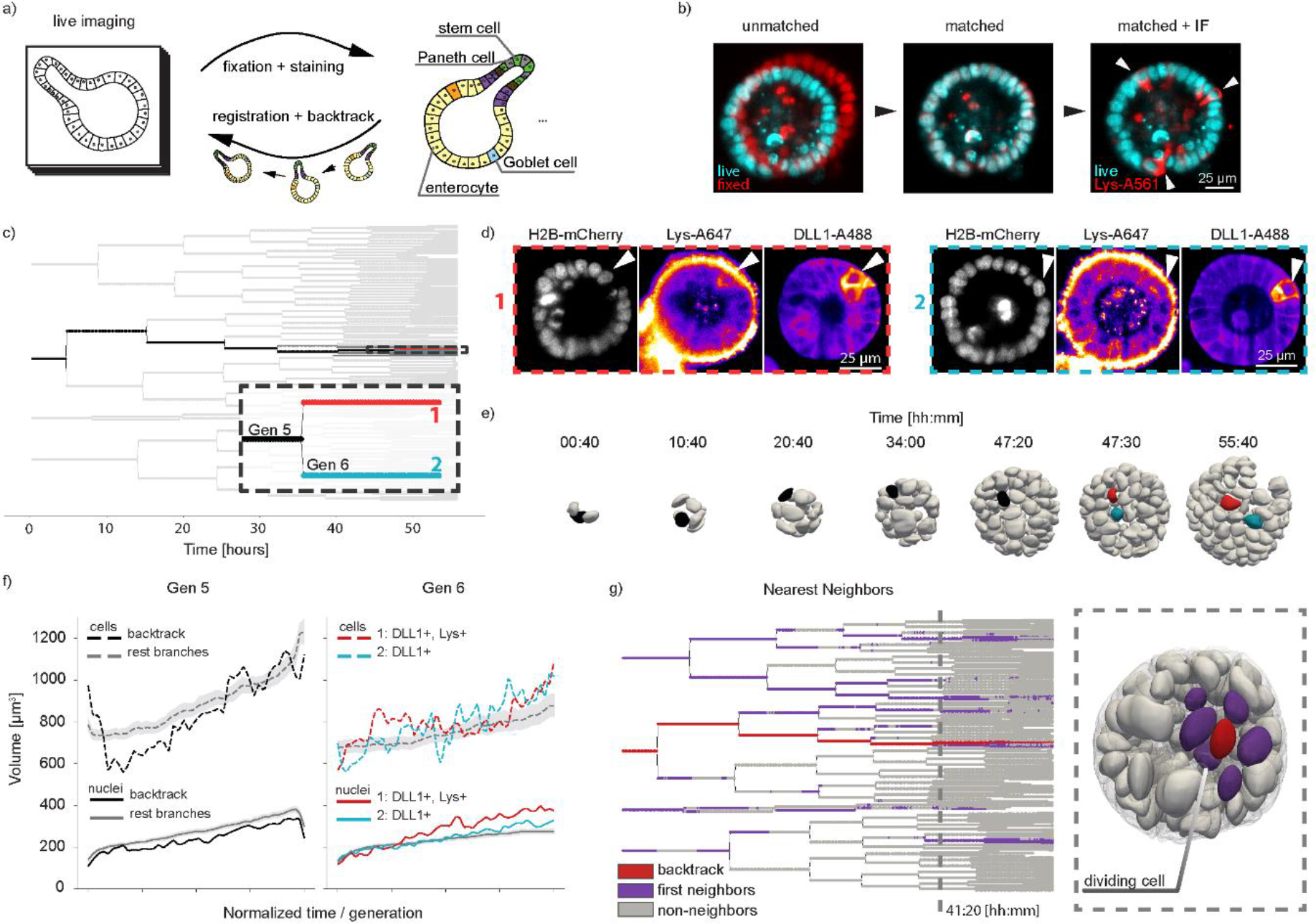
Fixation and backtracking after live-imaging. **a)** Fixation and backtracking strategy for light-sheet imaging of organoid growth (shown recording 007). **b)** After fixation a registration step may be needed to overlap the fixed nuclei with the nuclei as shown in the last timepoint of the timelapse. Left: midplane of raw data from last time-point H2B-mCherry recording (cyan) against same plane after fixation (red). Center: registration maps the fixed volume into the last time-point volume. Although overlap is not perfect, it is sufficient to maps each nucleus back. Right: After registration additional information from immunolabelling can be overlayed onto last time-point of the live recording, and so cells of interest can be backtracked. A total of 4 different experiments were performed with similar results. **c)** Lineage tree depicting the backtracking of two sister cells. **d)** After fixation, staining for DLL1 and Lysozyme shows two cells expressing these markers. Backtracking of them is shown in a).**e)** Nuclei volume distribution for the backtracked cells against all other cells per generation. **f)** Evaluation of nuclei and cell volumes for the backtracked cells against all other cells during generations 5 and 6, as depicted in c). For all other cells, the midline corresponds to the mean, whereas the gray region is the standard distribution. **g)** Nearest neighbor evaluation of the backtracked Lys+ cell (red). All nearest neighbors are depicting in magenta, with all remaining cells in gray. Dashed line on the lineage tree is presented as corresponding segmented meshes on the right. A dividing cell can be recognized by the interkinetic movement of its nucleus further apically. Source data are provided as a Source Data file.

In details, we fixed the sample at the end of the recording with 4% PFA, to then perform the immunolabelling protocol (see **Methods Section**). The pre-patterning of the FEP foil holding the sample was crucial, as without it the Matrigel drops were washed out. To account for organoid drifts, we imaged the entire fixation procedure, so that the organoids could be tracked during fixation, leading to a recovery of more than 80% (for more detailed information please see **Supplementary Note 6**). To register the fixed organoids to the last time-point of the live recording we used similarity transformations implemented in ITK and available via Elastix^28,29^ (used as a stand-alone tool, as exemplified in **Fig 4b**. For more information, please refer to the **Methods Section** and **Supplementary Note 6**). To test this approach, we imaged H2B-mCherry, mem9-GFP organoids until day 3 (**Supplementary Movie 3** left). It has been shown that between around day 1.5 intestinal organoids break symmetry through the appearance of the first differentiated cells of the secretory lineage (Paneth cells, Lysozyme). Preceding the appearance of Paneth cells there is the local establishment of a Notch-Delta lateral inhibition event, with future Paneth cells being typically Delta Like Ligand 1 positive (DLL1+)^6^. To analyze symmetry breaking, we fixed the organoids at 56 hours and stained for DLL1-Alexa488 and Lys-Alexa647 (**Fig 4c-e-, Supplementary Movie 3** right). Intriguingly, the two DLL1+ cells are two sister cells that were formed at the end of the division from generation 5 to generation 6, around 10 hours before fixation.

To follow cellular dynamics and changes of features of these specific cells in their spatial environment we analyzed nuclei and cell volumes (extracted with LSTree) per generation of the backtracked cells from generation 5 and 6 and compared them to all the other cells during the same generations (**Fig 4f**). Interestingly, cellular, and nuclear volumes of the backtracked cells do not seem to deviate relative to each other during generation 5. After cell division and entering generation 6, however, the nuclei volumes of both DLL1+ sister cells show an increased relative difference to one another, with the Lys+ cell having a slightly larger nucleus. Changes in nuclear volume related to appearance of DLL1+ signal was also observed in other datasets (**Supplementary Figure 6**).

Next, we evaluated the dynamics of neighbor exchange by cross-checking the closest cells to a backtracked cell(s) of interest at each time-point with LSTree. We examined if the progeny of these two sister cells had high level of mixing with other cells during cyst growth. From the visualization of the tracked neighbors on the lineage tree and segmented meshes (**Fig 4g**), it is apparent that neighbor exchanges, although distributed across the tree, do not happen often nor with many different cells, keeping an average of 5 cells.

The above results show that, by combining our light-sheet framework with standard fixation and registration techniques we can broaden the level of functional information, bridging it to the dynamical processes during live imaging. Consequently, we were capable of dissecting some initial dynamical elements preceding the formation of DLL1+ and Paneth cells in the context of the entire organoid development, analyzing the process of symmetry-breaking events across biological scales.

### Nuclei merging events during organoid growth

From our backtracking example it became apparent that one cell undergoes multiple rounds of failed divisions, with two daughter cells merging before a new division starts (**Fig 5a**). Upon further inspection of the other lineage trees, we realized that most of the datasets contained at least one merging event during early phase of organoid growth whereby two sister nuclei, at the end of their cell cycle, divided again into two instead of into four nuclei (**Supplementary Figure 7**). To investigate whether a failed division during the previous mitotic cycle was causing these nuclei merging, we examined the last step of the previous cell division. In all cases there was a problem during late cytokinesis, with the two sister cells never fully separating (**Fig 5a,b**). Nuclear volume for all the daughters arising after the merging event is clearly increased and cell volume followed the same behavior, roughly doubling in tetraploid cells (**Fig 5c,d**). To dismiss the possibility that these mitotic failures are caused by phototoxic effects of the imaging itself, we performed a time-course experiment with wild-type organoids grown from single cells under same medium conditions as the live recordings. We fixed organoids at days 2 and 3, staining them for e-Cadherin and DAPI. Despite the lack of continuous illumination, the resulting data showed many cysts with polynucleated cells, as well as cells with enlarged nuclei (**Fig 5e**).

**Figure 5:**
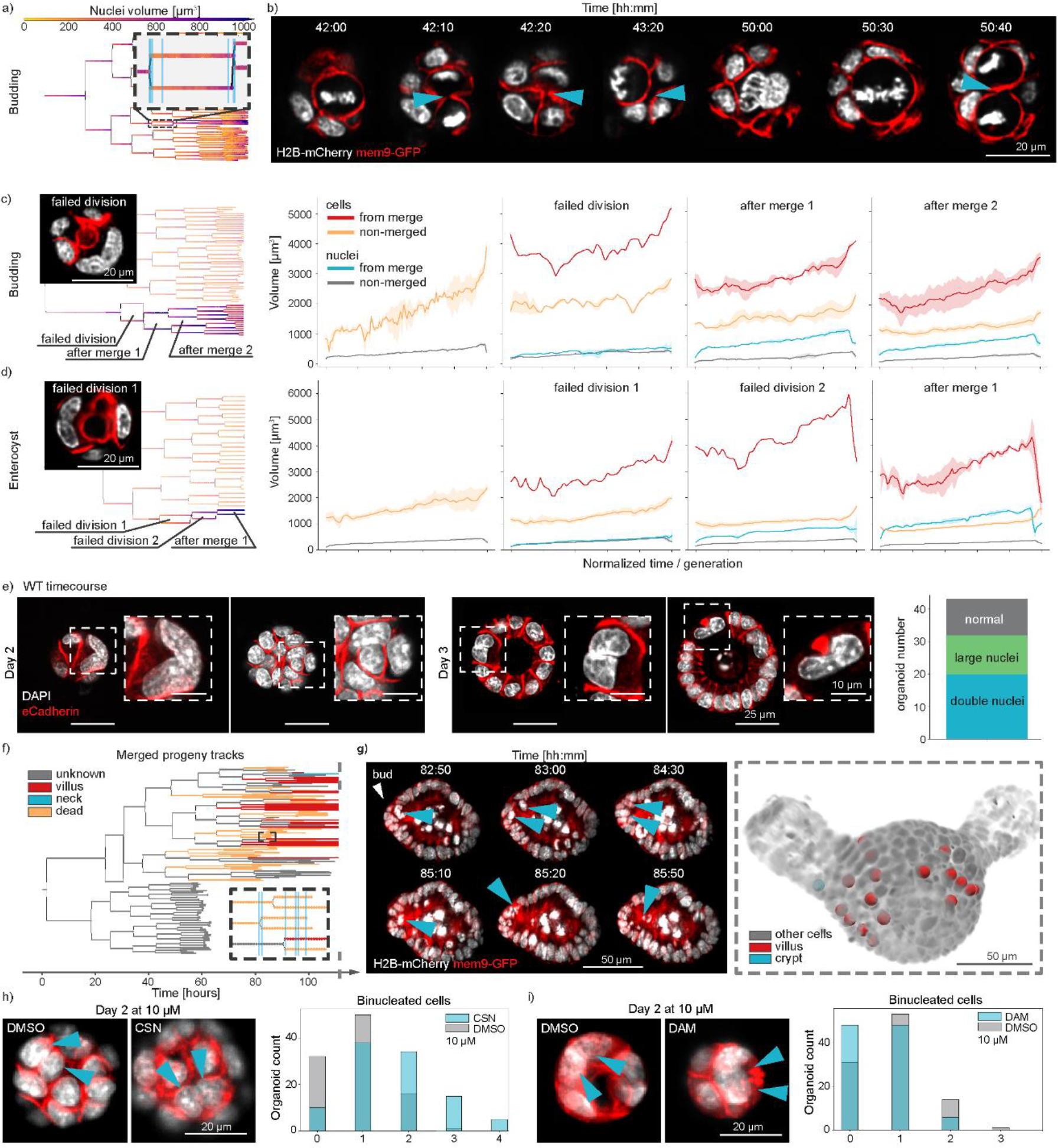
Merging events during early organoid growth. **a)** Example lineage tree with highlighted insert depicting a cell division where two nuclei divide again into two nuclei (recording 002). **b)** Still images of the light-sheet recording related to the dataset in a). Cyan arrows mark the position of a connecting region between the two nuclei until the following division occurs, where cytokinesis is successful. Corresponding locations for each depicted timepoint on the lineage tree are shown as cyan lines in the insert in a). **c**,**d)** Examples of different sequence of events following a failed division during budding organoid (recording 001) and enterocyst (recording 003) growth (left),with quantified nuclei and cell volumes for the highlighted events on the trees (right). Whenever more than one track is being evaluated for the same label, the full line represents the mean whereas the shaded region corresponds to 95% of the confidence interval. **e)** Timecourse data on wild-type intestinal organoids grown from single cells and fixed at days 2 and 3 after seeding. Staining with DAPI and e-Cadherin show binucleated cells and cells with large nuclei. Major axis length of all nuclei after a failed division (green) and all other cells (gray)shown on the right. **f)** Typical outcome of progeny from a failed division; *dead* tracks end with nuclei inside the lumen or basally extruded, unknown corresponds to tracks where the high level of cell packing and/or the low quality of the images made it impossible to continue and know their fate, and *alive* corresponds to tracks where the cells are still part of the epithelium until the end of the recording. **g)** Left: still images depicting timepoints highlighted in the inset in f), with nuclei ending in the lumen. Right: overlay of tracked cells from left panel onto the last time-point of the recording, showing the spatial organization of the cells in the crypt and vilus regions (recording 001). **h**,**i, left panel)** Example images of binucleated cells for both control and compound treated organoids, with nuclei highlighted via the arrows. **h**,**i, right panel)** Organoid count with specific number of binucleated cells for control (DMSO) and compound treated cases. 200 organoids have been randomly selected for evaluation on each case. Source data are provided as a Source Data file.

Using the framework we were able to follow the fate of the progeny of a cell that underwent cytokinesis failure and surprisingly we noticed that they can lead to cells that remain part of the epithelium until the end of the recordings without dying, when we can observe fully budded organoids or mature enterocysts (end of day 4) (**Fig 5f,g**). Yet, comparing to unaffected parts of the trees, this binucleation progeny typically has higher probability to be extruded into the lumen (∼46% for merged progeny against ∼5% for other cells). Another intriguing observation is that the remaining 54% of the cells are never localized to the crypt but to the villus (**Fig 5f, Supplementary Figure 8**). This is an interesting result, as it suggests that the cells that undergo cytokinesis failure, and might have chromosomal defect, do not migrate or differentiate into niche cells (Stem cells and Paneth cells) but stay as villus cells that are shorter lived. This might mean that there are mechanisms to maintain cellular integrity in the stem cell niche avoiding damaged cells in the crypt.

Molecularly, it is known that the Large tumor suppressor kinase 1 (Lats1) can influence cytokinesis failure via lack of inhibition of Lim kinase 1 (Limk1)^30-32^. This poses an interesting hypothesis on the role of mitotic failure in intestinal regeneration as Lats1 and Yap1 are master regulators of regenerative response of the intestinal epithelium^33,34^. Analysis of RNAseq from previous studies^6^, show decrease in Lats1 expression during initial days of organoid growth that mimic the regenerative response of the intestinal epithelium^6^. This regenerative response is achieved by downregulating Lats1 as a negative regulator of Yap1. We initially stained for Limk1 and could see cell-to-cell variability of its expression (**Supplementary Figure 9a**,**b**). To further analyze the role of Lats1 and Limk1 in the regulation of cytokinesis failure in the early days of organoid formation we perturbed Lats1 and Limk1 activity. Time-course imaging of Lats1 double knockouts^6^ showed several cysts with double nucleated cells that result from mitotic failures (**Supplementary Figure 9c**). Moreover, inhibition of Lats1 and Limk1 with chemical inhibitors (Truli and Damnacanthal, see **Methods Section** and **Supplementary Figure 9**), increased and decreased the number of bi-nucleated cells, respectively (**Fig 5h,i**). Lats1 inhibition also display an increase of Yap1 activation. This shows that in intestinal organoids formation, cytokinesis failure is regulated by Lats1 activity that in turn is a negative regulator of Limk1.

Taken together, through the multi scale approach of 3D segmentation, feature extraction and lineage tree analysis we were able to identify consistent polyploidy events during early intestinal organoid development and the fate of their progeny. Our framework allowed us to bridge the observed mitotic defects across scales towards the tissue scale, showing the end fate of the merged cells progeny and spatially locating them onto the mature organoid morphology. This is shedding new light on the robustness of a regenerative YAP cellular state, questioning the role of polyploidy in intestinal regeneration.

## Discussion

Here we have presented a unified light-sheet imaging framework tailored to intestinal organoid development. Our framework encompasses optimization of sample mounting, microscope recording and pre-processing, generating high-quality datasets with minimal storage needs. The image analysis and visualization part, named LSTree, is a comprehensive approach that uniquely combines image pre-processing, single cell tracking and multiscale segmentation and feature extraction along with a dedicated unified visualization and analysis tool. We show that this framework is capable of fully segmenting and tracking intestinal organoids as they grow from single cells for several days, bridging biological scales to the point when the organoid has hundreds of cells. Information on organoid, lumen, cell and nuclei volumes along with other multivariate features can be simultaneously visualized with the lineage tree data and further analyzed through a web-based Digital Organoid Viewer, facilitating a more global understanding of the dynamics acquired at subcellular resolution. The use of specialized neural networks dedicated to multiscale segmentation and lineage tree predictions allow the framework to be plastic enough to handle different kinds of live-imaging data as well as continuously improve through the utilization of dedicated tools for retraining of the models. Furthermore, the training architectures keep the number of parameters and computational resources to a minimum, relieving the needs for any highly specialized IT infrastructure, as they can be directly used on off-the-shelf workstations.

The combination of live imaging with standard fixation techniques, via the development of sample holder patterned with a cold stamp technique, allows the tracking of immunolabelled cells back in time and space and compare their features to all of the other cells over the entire organoid growth. Unlike lineage tracing in single cell RNA sequencing, which clusters cells by their RNA phenotypic fingerprint or a barcode, with our approach we can focus on the missing spatiotemporal organization in a causal way, as we follow the same cells over time. Consequently, we can now address the evolution of cells that can give rise to the first symmetry breaking event, keeping track of local interaction within the whole organoid.

Although suited to intestinal organoids, or to systems with smaller size, such as the mouse embryo, we recognize that particular challenges can be addressed to make our framework best suited to other case scenarios. For example, the application of our framework to larger organoids (surpassing diameters of 200 μm or being composed of highly dense cell aggregates) would best require light-sheet microscopes with more than one detection objective, or with sample rotation, so that the samples can be visualized from the opposite side as well, with the multi-view stacks properly fused afterwards. Furthermore, the utilization of multiphoton imaging could be of benefit to improve light-tissue penetration depth. Here the challenge is to provide multi-view imaging and yet keep the multi-sample aspect, so that systems with low growth efficiency can also be studied.

On the analysis side, the utilization of tracking spots for aiding in nuclei segmentation prediction shows to be an interesting approach to minimize the amount of hand annotated data for training. The extension of this approach towards cell volume estimation is a valid first approach, however, may still yield “noisy” cell volumes over time. To refine this, retraining with a few hand-annotated cells would be of good practice. Lastly, lineage tree prediction is still highly dependent on good temporal spacing and good nuclei segmentation. Especially for the challenging case of samples with low efficiency, such as the intestinal organoids, we tackle this issue with the careful adjustment of imaging parameters and of initial nuclei segmentation, as well as by allowing the workflow to receive input from other tracking methods such as Elephant. The possibility to use current state of the art tracking strategies such as Elephant or Mastodon helps especially to get first lineage trees done, and allows our LSTree workflow to be agnostic to only one approach. For the future we imagine that the inclusion of 3D ellipsoids instead of only spheres as weak annotation input for nuclei segmentation model training will aid in the nuclei segmentation quality, as the ellipsoids inherently carry more spatial information on the shape of the nuclei to be segmented.

Last, we present appearance of cells having cytokinesis-related mitotic errors leading to binucleation during early organoid growth. With LSTree we were able of rapidly verifying these errors across a multitude of different long-term recordings, showing that they are a consistent feature during early cyst growth. Interestingly, polyploidy has been associated with wound healing after injury^36^. In the liver, polyploid cells seem to have a tumor-suppressor role with polyploid cells occurring mostly due to cytokinesis failure and endoreduplication^37,38^. However, polyploid hepatic cells are mostly quiescent and do not divide unless the liver undergoes regenerative process due to a lesion. In contrast, our observed bi-nucleated cells do not undergo cell cycle arrest, but continue to divide for even multiple cycles, either with or without repeated clear mitotic failure. We also show that although these cells may appear in the crypt region during crypt formation (during days 3-4), they typically do not manage to reside in the crypt, whereas cells in the villus region remain part of the epithelium. Mitotic errors are negatively regulated by overactivation of Lats1 via regulation of Limk1. Since Lats has a direct implication during cytokinesis, we propose that during organoid growth and possibly intestinal regeneration there must be a balance between high-proliferative regenerative state – which is more error prone – and a counteracting checkpoint at the tissue scale to avoid mutations in the stem cell compartment. This way tissue integrity can be achieved fast, with any remaining mutations in the villus being eventually shed off via e.g. anoikis when cells reenter homeostasis.

In conclusion, with LSTree we can cross biological scales with unprecedented detail, as we can follow particular subcellular behaviors while keeping track on the entire tissue development over long periods of time. The usability of the tools presented can go far beyond the examples shown here, as they can be compatible with different light-sheet modalities making this framework also very useful in the study of other 3D cell cultures. In particular for the analysis, we believe LSTree to be a first step towards a comprehensive and quantitative framework dedicated to the creation of fully digital organoid maps, so that the intrinsic culture variabilities can be overcome with the creation of averaged organoids to be used as landmarks for future studies.

## Methods

### Ethics statement

All animal based studies have been approved by Basel Cantonal Veterinary Authorities and conducted in accordance with the Guide for Care and Use of Laboratory Animals.

### Organoid lines

Male and female outbred mice between 8 and 12 weeks old were used for all experiments. Regarding husbandry, all mice have a 12/12 hours day/night cycle. Medium temperature is kept at 22°C and relative humidity at 50%.

Mouse lines used for time-course experiments: C57BL/6 wild type (Charles River Laboratories), one 12 weeks old male and one 8 weeks female mice.

For all light-sheet movies, we used H2B-mCherry C57BL/6 x C3H F1 female intestines heterozygous for H2B-mCherry (received already as intestines, kind gift from T. Hiragi lab, EMBL). For H2B-mCherry/mem9-GFP organoids, H2B-mCherry organoids were infected with LV.EF1.AcGFP1-Mem-9 lentivirus particle (Clontech, Takara Bio USA). For the H2B-miRFP670 line, B6/N x R26 Fucci2 (Tg/+) intestines (kind gift from J. Skotheim lab, Stanford) were infected with pGK Dest H2B-miRFP670 (Addgene). For Lats DKO, Lats1Δ/Δ; Lats2Δ/Δ (LATS DKO, intestines as kind gift from Jeff Wrana, Department of Molecular Genetics, University of Toronto, Canada)^39^ time-course of published data^6^ was analyzed.

### Organoid culture

For initial organoid culture a section of the initial part of the small intestine was opened lengthwise and cleaned with cold PBS. Then, after removal of villi by scraping with a cold glass slide, the section was sliced into small fragments of roughly 2 mm in length. All fragments were then incubated in 2.5 mM EDTA/PBS at 4 °C for 30 min with shaking. Supernatant was removed and pieces of intestine were re-suspended in DMEM/F12 with 0.1% BSA. The tissue was then shaken vigorously. To collect the first fraction, the suspension was passed through a 70 μm strainer.

The remaining tissue pieces were collected from the strainer and fresh DMEM/F12 with 0.1% BSA was added, followed by vigorous shaking. The crypt fraction was again collected by passing through a 70 μm strainer. In total, 4 fractions were collected. Each fraction was centrifuged at 300g for 5 min at 4 °C. Supernatant was removed and the pellet was re-suspended into Matrigel with medium (1:1 ratio) and plated into 24 well plates. Organoids were kept in IntestiCult Organoid Growth Medium (STEMCELL Technologies) with 100 μg/ml Penicillin-Streptomycin for further amplification and maintenance.

### Organoid preparation for time-course experiments

WT organoids passage 10 were collected 5-7 days after passaging and digested with TrypLE (Thermo Fisher Scientific) for 20 min at 37 °C. The resulting dissociated cells were filtered through a 30 µm cell strainer (Sysmex) and single alive cells were sorted by FACS (Becton Dickinson Influx cell sorter with BD FACS Sortware 1.2.0.142, or Becton Dickinson FACSAria III using BD FACSDiva Software Version 8.0.1). Forward scatter and side scatter properties were used to remove cell doublets and dead cells. The collected cells were resuspended in ENR medium (advanced DMEM/F-12 with 15 mM HEPES (STEM CELL Technologies) supplemented with 100 μg/ml Penicillin-Streptomycin, 1×Glutamax (Thermo Fisher Scientific), 1×B27 (Thermo Fisher Scientific), 1xN2 (Thermo Fisher Scientific), 1mM N-acetylcysteine (Sigma), 500ng/ml R-Spondin (kind gift from Novartis), 100 ng/ml Noggin (PeproTech) and 100 ng/ml murine EGF (R&D Systems) and mixed 1:1 with Matrigel (Corning). Cells were seeded at a density of 3000 cells per 5ul drops per well of 96 well imaging plates (Greiner, 655090)(2 plates and 3 wells per condition). After 20 min of solidification at 37 °C, 100 µl of medium was overlaid. From day 0 to day 1, ENR was supplemented with 20% Wnt3a-conditioned medium (Wnt3a-CM), 10 μM Y-27632 (ROCK inhibitor, STEMCELL Technologies) and 3 µM of CHIR99021 (GSK3B inhibitor, STEMCELL Technologies, cat # 72054). From day 1 to 3 ENR was supplemented with 20% Wnt3a-CM and 10 μM Y-27632.

### Fixed sample preparation and time-course imaging

Organoids are embedded in a Matrigel droplet. Due to the nature of the droplet, individual organoids are located at different heights in the Matrigel drop. To allow imaging of all organoids within a similar z-range, each 96-well plate was centrifuged at 847 g for 10 min in a pre-cooled centrifuge at 10 °C prior to fixation. Organoids were fixed in 4% PFA (Electron Microscopy Sciences) in PBS for 45 min at room temperature. For time course and compound experiments, organoids were permeabilized with 0.5% Triton X-100 (Sigma-Aldrich) for 1 h and blocked with 3% Donkey Serum (Sigma-Aldrich) in PBS with 0.1% Triton X-100 for 1 h.

### WT imaging

For the images in Figure 5e, membrane staining with E-Cadherin (BD Biosciences, # 610182) was done at 1:300 ratio in Blocking buffer for 20 hours at 4°C. DAPI staining was performed at concentration of 300 nM for 30 min at room temperature. All secondary antibodies were added at 1:300 for 1 hour in room temperature. Cell nuclei were stained with 20 μg/ml DAPI (4’,6-Diamidino-2-Phenylindole, Invitrogen) in PBS for 15 min. High-throughput imaging was done with an automated spinning disk microscope from Yokogawa (CellVoyager 7000S), with an enhanced CSU-W1 spinning disk (Microlens-enhanced dual Nipkow disk confocal scanner), a 40x (NA = 0.95) Olympus objective, and a Neo sCMOS camera (Andor, 2,560 × 2,160 pixels). For imaging, an intelligent imaging approach was used in the Yokogawa CV7000 (Search First module of Wako software). For each well, one field was acquired with 2x resolution in order to cover the complete well. This overview fields were then used to segment individual organoids on the fly with a custom written ImageJ macro which outputs coordinates of individual organoid positions. These coordinated were then subsequently imaged with high resolution (40x, NA = 0.95). For each site, z-planes spanning a range up to 140 μm were acquired. For the data in Figure 5e,h and in Supplementary Figure 9 2 μm z-steps were used.

### Lats-DKO

Analysed data stems from a previous publication^6^, with Lats DKO organoids dissociated into single cells and plated into 96 well plates, fixed and stained with DAPI following the published protocols. Tamoxifen induction (1:1000) was kept in the medium until fixation time.

### RXRi

RXR inhibition was achieved by adding the Cpd2170 RXR antagonist^7^ compound at 1:2000 ratio to the medium from the moment single cells were seeded onto the light-sheet holder. The compound was kept throughout the data acquisition. Organoids used for this experiment had been infected with H2B-iRFP670 for live nuclei labeling.

### Inhibition experiments: Lats1/2 and Limk1 inhibition time-course

For the evaluation of binucleated cells in Fig5h-i and Supplementary Figure 9, FACS sorted (Becton Dickinson Influx cell sorter with BD FACS Sortware 1.2.0.142, or Becton Dickinson FACSAria III using BD FACSDiva Software Version 8.0.1). WT mouse intestinal organoids at passage 10 were dissociated and grown from single cells as described above. Inhibitors were resuspended in DMSO and serially diluted in medium to their final working concentration and added on day 0 (Lats1/2 inhibitor Truli^40^ (CSNpharm, # CSN26140) or the Limk1 inhibitor Damnacanthal^41^ (Tocris, # 1936)). One plate was fixed with 4% PFA on day 2 (48hrs after plating) and the other one on day 3 (72hrs after plating) as described in the previous section. At the end of the time course all plates were permeabilized with 0.5% Triton X-100 (Sigma-Aldrich) for 1 h and blocked with 3% Donkey Serum (Sigma-Aldrich) in PBS with 0.1% Triton X-100 for 1 h. Primary antibodies were diluted in blocking as follow: anti-e-Cadherin (BD Biosciences, # 610182) 1:400, anti-Limk1 (Abcam, # ab194798) 1:400 and anti-Yap1 (Cell Signaling, # 14074) 1:400 and incubated for 1h at RT on a shaking plate. The primary antibodies were washed with PBS 3×10min at RT on a shaking plate. Both secondary antibodies (Alexa Fluor 568 donkey anti mouse, Thermo Fisher Scientific; A10042 and Alexa Fluor 488 donkey anti rabbit, Thermo Fisher Scientific; A-21202) were diluted 1:400 and incubated for 2hrs at RT on a shaking plate. The plates were then washed with PBS 3×10min at RT on a shaking plate and cell nuclei were stained with 20 μg/ml DAPI (4’,6-Diamidino-2-Phenylindole, Invitrogen) in PBS for 15 min. Plates were then covered in aluminum foil and imaged with the ImageXpress from MolecularDevices. Stacks were acquired with 20X objective (0.3417 μm in X and Y) and 3 μm steps. For analysis, 200 randomly picked organoids were selected for each condition and the number of binucleated cells present on each one evaluated.

### Light-sheet sample preparation

H2b-mCherry / mem9-GFP and H2B-iRFP670 organoids were collected and digested with TrypLE (Thermo Fisher Scientific) for 20 min at 37 °C. Alive double positive (mCherry/GFP) cells were sorted by FACS (Becton Dickinson Influx cell sorter with BD FACS Sortware 1.2.0.142, or Becton Dickinson FACSAria III using BD FACSDiva Software Version 8.0.1). and collected in medium containing advanced DMEM/F-12 with 15 mM HEPES (STEM CELL Technologies) supplemented with 100 μg/ml Penicillin-Streptomycin, 1×Glutamax (Thermo Fisher Scientific), 1×B27 (Thermo Fisher Scientific), 1xN2 (Thermo Fisher Scientific), 1mM N-acetylcysteine (Sigma), 500ng/ml R-Spondin (kind gift from Novartis), 100 ng/ml Noggin (PeproTech) and 100 ng/ml murine EGF (R&D Systems). 2500 cells were then embedded in 5ul drop of Matrigel/medium in 60/40 ratio. Drops were placed in the imaging chamber and incubated for 20 min before being covered with 1ml of medium. For the first three days, medium was supplemented with 20% Wnt3a-CM and 10 μM Y-27632 (ROCK inhibitor, STEMCELL Technologies). For the first day, in addition, 3μM of CHIR99021 (STEMCELL Technologies) were supplemented. After 2 hours incubation in a cell culture incubator the imaging chamber was transferred to the microscope kept at 37C and 5% CO2.

### Light-sheet imaging

For all light-sheet experiments a LS1-Live dual illumination and inverted detection microscope from Viventis Microscopy Sàrl was used. Different single cells were selected as starting positions and imaged every 10 min for up to 5 days. A volume of 150 -200μm was acquired with a Z spacing of 2μm between slices and 100 ms exposure time for each slice. Laser intensity was kept to a minimum necessary to still obtain reasonable signal to noise from the raw data, while keeping phototoxicity to a minimum possible. Medium was exchanged manually under the microscopy every day.

### Fixation on time-lapse recordings

Organoids are embedded in 5 μm Matrigel droplets which are deposited at equal distances on top of the FEP foil of the light-sheet sample holder. After live imaging is done, the medium is replaced by 4% PFA in PBS, and left in the chamber for maximum 30 minutes at 37°C in the microscope. After fixation the organoids were permeabilized with 0.5% Triton X-100 (Sigma-Aldrich) for 1 h and blocked with 3% Donkey Serum (Sigma-Aldrich) in PBS with 0.1% Triton X-100 for 1 h. For the images in Figure 4, the cyst was stained with DLL1 antibody (R&D Systems, # AF3970) at 1:100 ratio and left overnight at 4°C. For Lysozyme (Dako, # A0099) we used a 1:400 ratio for 3 hours at room temperature.

### Registration for back-tracking after fixation of time-lapses

Since PFA fixation causes the Matrigel droplet to flatten, we perform imaging while fixation is taking place. Typically we observe no change within the first 5 minutes, whereas after that there is a sudden increase in organoid movement towards the bottom of the sample holder. To take this into account, we increased the imaging volume and step size to be able to encompass a larger volume and still track the organoid. For the data in Figure 4 we increased stack size from 150 μm and 2 μm step size to 300 μm at 3 μm step size. However, larger values can also be used.

Nonetheless, the flattening of the droplet will lead the organoids to rotate or translate in space. Furthermore, PFA fixation also changes the shape of tissue samples by shrinking or swelling. To bridge the translational, rotational and rescaling of the organoids during fixation procedures, we registered fixed organoids using Elastix v4.900 (https://elastix.lumc.nl/). Since Elastix can be directly installed from the repository as pre-compiled libraries, we refrained from embedding the registration into LSTree, and left it as a stand-alone tool. For all registrations using the similarity transform, a base parameter file set for performing similarity transformations was used and eventually modified so that best results could be achieved. An example of the registration parameters is provided in ‘Elastix_parameter_Affine.txt’ file in the Supplementary Software.

### LSTree modules

LSTree (https://github.com/fmi-basel/LSTree) is a luigi-based workflow (https://github.com/spotify/luigi) which encompasses jupyter notebooks for cropping and general utilities, as well as luigi lasks for denoising, deconvolution and multiscale segmentation and tree-prediction, along with feature extraction. Pre-processing steps rely mostly on cropping and registration, denoising, and deconvolution steps. Deconvolution was based on flowdec (https://github.com/hammerlab/flowdec). Although not part of LSTree itself, improvements in the microscope software (on-the-fly LZW compression, position dependent illumination alignment) were performed in collaboration with Viventis Microscopy Sàrl and are now part of their current microscope software. A lzw compression python code (‘parallel_image_compressor.py’) is available in the **Supplementary Software**.

Detailed information regarding pre-processing, segmentation strategies and feature extraction can be found in **Supplementary Text**.

### Software

For deconvolution of the images, PSFs were averaged using the PSF Distiller from Huygens compute engine 20.10.1p1. For visualization of images ImageJ v.1.53h and Paraview 5.8.0 were used, and Elastix v4.900 was used for registration of organoids.

### IT requirements

The LSTree analysis tasks have been trained and used on a workstation with following specifications: 16 core Intel Xeon W-2145, 64 GB 2666MHz DDR4 RAM equipped with a Nvidia Quadro RTX 6000 GPU with 24 GB VRAM and using Ubuntu 18.04.6 LTS. All code runs with Nvidia cudatoolkit 10.1, and cuDNN 7.

Minimally, one would need 16 GB of RAM and a Tensorflow compatible GPU with at least 8 GB of VRAM. Since many of the steps of the pipeline run in parallel, a higher number of CPUs is also desirable.

A step-by-step guide on installation and on how to run the example data provided can be found in the repository (www.github.com/fmi-basel/LSTree).

### Statistics & Reproducibility

For all experiments no statistical method was used to predetermine sample size. Sample size was determined based on previous related studies in the field^11,16,27^. For long-term live imaging experiments, we assumed that the amount of timepoints comprised in the 7 different datasets would be sufficient to test the framework. In addition, 12 other datasets from previous publication^27^ were used for further challenging the analysis framework). No data were excluded from the analyses. Samples were randomly assigned. Investigators were not blinded to allocation during experiments and outcome assessment.

## Supporting information

Supplementary Software

## Data Availability

Source data are provided with this paper. A minimum example to test LSTree is provided within the repository. The light-sheet data and time-course data generated in this study have been deposited in the Zenodo database under accession code 10.5281/zenodo.6828906 [https://zenodo.org/record/6828906]^42^. Due to storage space restrictions, for source light-sheet image data please contact Prisca Liberali for more information.

## Code Availability

LSTree can be found publicly in GitHub (https://github.com/fmi-basel/LSTree) with its latest release referenced also in Zenodo (DOI: 10.5281/zenodo.6826914)^17^. All other code used in this work is present in the Supplementary Software.

## Acknowledgements

We thank M. Rempfler for helpful discussions on framework implementation and support, D. Vischi, E. Tagliavini and Sjoerd van Eeden for IT support, T.-O. Buchholz for aid with segmentation evaluation and neural network support, J. Eglinger for help with KNIME workflow, Q. Yang for discussions on pre-processing, S. Xie for helpful discussions on biological aspects, A. Peters for light-sheet, H. Kohler for sorting, P. Argast for help with building the FEP foil aluminum stamp, C. Tsiairis, L. Gelman and laboratory members for reading the manuscript. Funding: EMBO (ALTF 571-2018 to G.M.), SNSF (POOP3_157531 to P.L.). This work received funding from the ERC under the European Union’s Horizon 2020 research and innovation programme (grant agreement no. 758617).

## Author contributions

P.L. and G.M. conceived and P.L. supervised the study, P.L., G.M., and A.B. designed the experiments, F.Ma., L.C.M. and G.M. cultured the organoids, G.M. and A.B. recorded the time-lapses, N.R. and G.M performed backtracking experiments. F.Mo. trained and evaluated tracking predictions with Elephant. P.S. wrote and implemented compression and position dependent illumination code into microscope software, A.B. created first Mastodon trees, R.O. wrote first LSTree workflow with the support of G.M., G.M. and L.C.M. performed time-course experiments. G.M., R.O. and P.L. analysed the data from the time-lapses, G.M, L.C.M. and P.L. analysed the time-course experiments, G.M., R.O. and P.L. wrote the paper.

Correspondence should be sent to Prisca Liberali.

