## Supplementary Software for "Multiscale light-sheet organoid imaging framework": Elastix_parameters_Affine.txt

// Example parameter file for affine registration
// C-style comments: //

// The internal pixel type, used for internal computations
// Leave to float in general. 
// NB: this is not the type of the input images! The pixel 
// type of the input images is automatically read from the 
// images themselves.
// This setting can be changed to "short" to save some memory
// in case of very large 3D images.
(FixedInternalImagePixelType "float")
(MovingInternalImagePixelType "float")

// The dimensions of the fixed and moving image
// Up to elastix 4.5 this had to be specified by the user.
// From elastix 4.6, this is not necessary anymore.
//(FixedImageDimension 2)
//(MovingImageDimension 2)

// Specify whether you want to take into account the so-called
// direction cosines of the images. Recommended: true.
// In some cases, the direction cosines of the image are corrupt,
// due to image format conversions for example. In that case, you 
// may want to set this option to "false".
(UseDirectionCosines "true")

// **************** Main Components **************************

// The following components should usually be left as they are:
(Registration "MultiResolutionRegistration")
(Interpolator "BSplineInterpolator")
(ResampleInterpolator "FinalBSplineInterpolator")
(Resampler "DefaultResampler")

// These may be changed to Fixed/MovingSmoothingImagePyramid.
// See the manual.
(FixedImagePyramid "FixedRecursiveImagePyramid")
(MovingImagePyramid "MovingRecursiveImagePyramid")

// The following components are most important:
// The optimizer AdaptiveStochasticGradientDescent (ASGD) works
// quite ok in general. The Transform and Metric are important
// and need to be chosen careful for each application. See manual.
// IMPORTANT: the metric will be determining how the effective the uniform scaling is applied
(Optimizer "AdaptiveStochasticGradientDescent")
(Transform "SimilarityTransform")
//(Metric "AdvancedMattesMutualInformation")
(Metric "AdvancedNormalizedCorrelation")
// ***************** Transformation **************************

// Scales the affine matrix elements compared to the translations, to make
// sure they are in the same range. In general, it's best to  
// use automatic scales estimation:
(AutomaticScalesEstimation "true")

// Automatically guess an initial translation by aligning the
// geometric centers of the fixed and moving.
(AutomaticTransformInitialization "true")

// Whether transforms are combined by composition or by addition.
// In generally, Compose is the best option in most cases.
// It does not influence the results very much.
(HowToCombineTransforms "Compose")

// ******************* Similarity measure *********************

// Number of grey level bins in each resolution level,
// for the mutual information. 16 or 32 usually works fine.
// You could also employ a hierarchical strategy:
//(NumberOfHistogramBins 16 32 64)
(NumberOfHistogramBins 256)

// If you use a mask, this option is important. 
// If the mask serves as region of interest, set it to false.
// If the mask indicates which pixels are valid, then set it to true.
// If you do not use a mask, the option doesn't matter.
(ErodeMask "false")

// ******************** Multiresolution **********************

// The number of resolutions. 1 Is only enough if the expected
// deformations are small. 3 or 4 mostly works fine. For large
// images and large deformations, 5 or 6 may even be useful.
(NumberOfResolutions 4)

// The downsampling/blurring factors for the image pyramids.
// By default, the images are downsampled by a factor of 2
// compared to the next resolution.
// So, in 2D, with 4 resolutions, the following schedule is used:
//(ImagePyramidSchedule 8 8  4 4  2 2  1 1 )
// And in 3D:
//(ImagePyramidSchedule 8 8 8  4 4 4  2 2 2  1 1 1 )
// You can specify any schedule, for example:
//(ImagePyramidSchedule 4 4  4 3  2 1  1 1 )
// Make sure that the number of elements equals the number
// of resolutions times the image dimension.

// ******************* Optimizer ****************************

// Maximum number of iterations in each resolution level:
// 200-500 works usually fine for affine registration.
// For more robustness, you may increase this to 1000-2000.
(MaximumNumberOfIterations 500)

// The step size of the optimizer, in mm. By default the voxel size is used.
// which usually works well. In case of unusual high-resolution images
// (eg histology) it is necessary to increase this value a bit, to the size
// of the "smallest visible structure" in the image:
//(MaximumStepLength 1.0)

// **************** Image sampling **********************

// Number of spatial samples used to compute the mutual
// information (and its derivative) in each iteration.
// With an AdaptiveStochasticGradientDescent optimizer,
// in combination with the two options below, around 2000
// samples may already suffice.
(NumberOfSpatialSamples 2048)

// Refresh these spatial samples in every iteration, and select
// them randomly. See the manual for information on other sampling
// strategies.
(NewSamplesEveryIteration "true")
(ImageSampler "Random")

// ************* Interpolation and Resampling ****************

// Order of B-Spline interpolation used during registration/optimisation.
// It may improve accuracy if you set this to 3. Never use 0.
// An order of 1 gives linear interpolation. This is in most 
// applications a good choice.
(BSplineInterpolationOrder 1)

// Order of B-Spline interpolation used for applying the final
// deformation.
// 3 gives good accuracy; recommended in most cases.
// 1 gives worse accuracy (linear interpolation)
// 0 gives worst accuracy, but is appropriate for binary images
// (masks, segmentations); equivalent to nearest neighbor interpolation.
(FinalBSplineInterpolationOrder 3)

//Default pixel value for pixels that come from outside the picture:
(DefaultPixelValue 0)

// Choose whether to generate the deformed moving image.
// You can save some time by setting this to false, if you are
// only interested in the final (nonrigidly) deformed moving image
// for example.
(WriteResultImage "true")

// The pixel type and format of the resulting deformed moving image
(ResultImagePixelType "float")
(ResultImageFormat "mhd")
