## Supplementary Software for "Multiscale light-sheet organoid imaging framework": mouse_config.cfg

```
# General ##############################################################
[DEFAULT]
# pattern=/tungstenfs/scratch/gliberal/Users/quingust/Manuscripts/TechnicalPaper/EllenbergLab-MouseEmbryos/Strnad_etal_tracking_data/registered_images_scaled/TestDataset/{subdir}/{fname}-T{time:04d}.{ext}
pattern={subdir}/{fname}-T{time:04d}.{ext}

# set the same default number of threads of all multithreaded tasks
# when processing a single dataset use n_threads ~= number of cores
# Note that some tasks actually use multiprocessing when internal operations do not release the GIL
n_threads=8

[resources]
gpu=1
pool_workers=16
memory=64000

[core]
workers=16
log_level=INFO
outdir = /tungstenfs/scratch/gliberal/Users/quingust/Manuscripts/TechnicalPaper/EllenbergLab-MouseEmbryos/Strnad_etal_tracking_data/registered_images_scaled/TrainingSets/

# Denoising/Deconvolution ##############################################
[BuildDenoiseTrainingRecordTask]
training_base_dir=/tungstenfs/scratch/gliberal/Users/ortiraph/datasets/LSTree_training/denoise
base_dir=/tungstenfs/scratch/gliberal/Users/quingust/Manuscripts/TechnicalPaper/EllenbergLab-MouseEmbryos/Strnad_etal_tracking_data/registered_images_scaled/TrainingSets/

n_images=30
train_fraction=0.9
valid_fraction=0.1
min_patch_size=(512,512)

[DenoiseTrainingTask]
training_base_dir=/tungstenfs/scratch/gliberal/Users/ortiraph/datasets/LSTree_training/denoise
base_dir=/tungstenfs/scratch/gliberal/Users/quingust/Manuscripts/TechnicalPaper/EllenbergLab-MouseEmbryos/Strnad_etal_tracking_data/registered_images_scaled/TrainingSets/
# images_dirs=["*/EGFP", "*/mCherry"]
images_dirs=["*/mCherry"]

downsampling_factor=(4,)
n_downsampling_channels=16
n_groups=8
dilation_rates=(1, 2, 4)
channels_per_group=32
n_steps=5
dropout=0.1

train_batch_size=16
valid_batch_size=32
epochs=200
n_restarts=5
learning_rate=0.0001
patch_size=(128,128,1)
suffix=20200311

#resume_weights=PATH_TO_CURRENT_MODEL/weights_latest.h5

intensity_offset_sigma=0.5
intensity_scaling_bounds=(0.1, 10.)

[ChannelBoundsTask]


[DenoiseTask]
out_suffix=-Denoised

[DeconvolutionTask]
psf_dir=/tungstenfs/scratch/gliberal/Users/ortiraph/datasets/LSTree_training/deconv/20190830
out_suffix=-Deconv
niter=128
#~max_patch_size=(512,512,512)
max_patch_size=(9999,9999,9999)

[MultiDeconvolutionTask]
# ch_subdirs=["EGFP", "mCherry"]
ch_subdirs=["mCherry"]
movie_dirs=["/tungstenfs/scratch/gliberal/Users/quingust/Manuscripts/TechnicalPaper/EllenbergLab-MouseEmbryos/Strnad_etal_tracking_data/registered_images_scaled/TrainingSets/*-*"]

# Lineage ##############################################################
[TreePropsTask]
out_subdir=tree_props
xml_tree=embryo_mamut.xml

# Nuclei segmentation ##################################################
[BuildNucleiTrainingRecordTask]
training_base_dir=/tungstenfs/scratch/gliberal/Users/quingust/Manuscripts/TechnicalPaper/EllenbergLab-MouseEmbryos/Strnad_etal_tracking_data/registered_images_scaled/trainingModels/nuclei_seg/969696_raw
annot_subdir=nuclei_annot
ch_subdir = mCherry
spacing=(0.52, 0.52, 0.52)
train_fraction=0.9 
valid_fraction=0.1
min_patch_size=(96,96,96)
patch_margins=(40, 40, 40)

[BuildWeakNucleiTrainingRecordTask]
training_base_dir=/tungstenfs/scratch/gliberal/Users/quingust/Manuscripts/TechnicalPaper/EllenbergLab-MouseEmbryos/Strnad_etal_tracking_data/registered_images_scaled/trainingModels/nuclei_seg/969696_raw
spacing=(0.52, 0.52, 0.52)
ch_subdir = mCherry
train_fraction=0.9
valid_fraction=0.1
## CHANEG TO AVERAGE SIZE OF NUCLEI
min_patch_size=(96,96,96)
patch_margins=(40,40,40)

[NucleiWeakAnnotTask]
ch_subdir=mCherry
out_subdir=nuclei_weak_annot
movie_dirs=["/tungstenfs/scratch/gliberal/Users/quingust/Manuscripts/TechnicalPaper/EllenbergLab-MouseEmbryos/Strnad_etal_tracking_data/registered_images_scaled/TrainingSets/*-*"]

[NucleiSegmentationTrainingTask]
training_base_dir=/tungstenfs/scratch/gliberal/Users/quingust/Manuscripts/TechnicalPaper/EllenbergLab-MouseEmbryos/Strnad_etal_tracking_data/registered_images_scaled/trainingModels/nuclei_seg/969696_raw
movie_dirs=["/tungstenfs/scratch/gliberal/Users/quingust/Manuscripts/TechnicalPaper/EllenbergLab-MouseEmbryos/Strnad_etal_tracking_data/registered_images_scaled/TrainingSets/*-*"]

downsampling_factor=(8,8,8)
n_downsampling_channels=32
n_groups=4
dilation_rates=(1, 2, 4, 8)
channels_per_group=32
n_steps=5
dropout=0.1
n_classes=2
spacing=(0.52, 0.52, 0.52)

train_batch_size=4
train_batches_per_epoch=200
valid_batch_size=8
epochs=300
n_restarts=2
learning_rate=0.0001
patch_size=(96,96,96,1)
suffix=20220510-scratch_raw_2

intra_margin=2.0
inter_margin=6.0
jaccard_hinge=0.3
jaccard_eps=0.1

# resume_weights=/tungstenfs/scratch/gliberal/Users/quingust/Manuscripts/TechnicalPaper/EllenbergLab-MouseEmbryos/Strnad_etal_tracking_data/registered_images_scaled/trainingModels/nuclei_seg/OLD/out/RDCNet-F8-DC32-OC5-G4-DR1-2-4-8-GC32-S5-D0.1_20220222_w_validation/weights_latest.h5

# resume_weights=/tungstenfs/scratch/gliberal/Users/quingust/Manuscripts/TechnicalPaper/EllenbergLab-MouseEmbryos/Strnad_etal_tracking_data/registered_images_scaled/trainingModels/nuclei_seg/969696_raw/out/RDCNet-F8-DC32-OC5-G4-DR1-2-4-8-GC32-S5-D0.1_20220509-scratch_raw/weights_latest.h5

intensity_offset_sigma=0.5
intensity_scaling_bounds=(0.1, 10.)

[NucleiSegmentationTask]
ch_subdir=mCherry
out_subdir=nuclei_segmentation-20220510-scratch_raw_2

#if first time, predict nuclei from the datasets where trees existed and were used for training a nuclei segmentation model from scratch
[MultiNucleiSegmentationTask]
movie_dirs=[
    "/tungstenfs/scratch/gliberal/Users/quingust/Manuscripts/TechnicalPaper/EllenbergLab-MouseEmbryos/Strnad_etal_tracking_data/registered_images_scaled/TrainingSets/*-*",
    # "/tungstenfs/scratch/gliberal/Users/quingust/Manuscripts/TechnicalPaper/EllenbergLab-MouseEmbryos/Strnad_etal_tracking_data/registered_images_scaled/TrainingSets/006-*",
    "/tungstenfs/scratch/gliberal/Users/quingust/Manuscripts/TechnicalPaper/EllenbergLab-MouseEmbryos/Strnad_etal_tracking_data/registered_images_scaled/TestSets/*-*"
    ]

# Cell segmentation ##################################################
[BuildLumenTrainingRecordTask]
training_base_dir=/tungstenfs/scratch/gliberal/Users/ortiraph/datasets/LSTree_training/cell_seg
ch_subdir=EGFP
annot_subdir=lumen_annot
spacing=(0.52,0.52,0.52)
train_fraction=0.9
valid_fraction=0.1
min_patch_size=(256,256,256)
patch_margins=(80,80,80)

[BuildWeakCellTrainingRecordTask]
training_base_dir=/tungstenfs/scratch/gliberal/Users/ortiraph/datasets/LSTree_training/cell_seg
ch_subdir=EGFP
spacing=(0.52,0.52,0.52)
train_fraction=0.9
valid_fraction=0.1
min_patch_size=(256,256,256)
patch_margins=(40,40,40)

[CellSegmentationTrainingTask]
training_base_dir=/tungstenfs/scratch/gliberal/Users/ortiraph/datasets/LSTree_training/cell_seg
movie_dirs_lumen=["/tungstenfs/scratch/gliberal/Users/ortiraph/datasets/LSTree_source/*-*"]
movie_dirs_cell=["/tungstenfs/scratch/gliberal/Users/ortiraph/datasets/LSTree_source/000-*",
                 "/tungstenfs/scratch/gliberal/Users/ortiraph/datasets/LSTree_source/001-*",
                 "/tungstenfs/scratch/gliberal/Users/ortiraph/datasets/LSTree_source/002-*",
                 "/tungstenfs/scratch/gliberal/Users/ortiraph/datasets/LSTree_source/003-*",
                 "/tungstenfs/scratch/gliberal/Users/ortiraph/datasets/LSTree_source/006-*",
                 "/tungstenfs/scratch/gliberal/Users/ortiraph/datasets/LSTree_source/007-*",
                 "/tungstenfs/scratch/gliberal/Users/ortiraph/datasets/LSTree_source/009-*"]

downsampling_factor=(1,8,8)
n_downsampling_channels=32
n_groups=4
dilation_rates=(1, 2, 4, 8)
channels_per_group=32
n_steps=5
dropout=0.1
n_classes=3
spacing=(0.52,0.52,0.52)

train_batch_size=8
train_batches_per_epoch=200
valid_batch_size=32
epochs=200
n_restarts=5
learning_rate=0.0001
patch_size=(24,192,192,1)
suffix=20210227

intra_margin=2.0
inter_margin=6.0
jaccard_hinge=0.3
jaccard_eps=1.0

#resume_weights=PATH_TO_CURRENT_MODEL/weights_latest.h5

intensity_offset_sigma=0.5
intensity_scaling_bounds=(0.1, 10.)

[CellSegmentationTask]
ch_subdir=EGFP
out_subdir_lumen=lumen_segmentation
out_subdir_cell=cell_segmentation

[MultiCellSegmentationTask]
movie_dirs=["/tungstenfs/scratch/gliberal/Users/quingust/Manuscripts/TechnicalPaper/EllenbergLab-MouseEmbryos/Strnad_etal_tracking_data/registered_images_scaled/TrainingSets/017-*"]

# Features #############################################################
[ExtractFeaturesTask]
out_subdir=features-20220511-scratch_raw_3
nuclei_subdir=mCherry

[MultiAggregateFeaturesTask]
# datasets to process that are tracked
movie_dirs=["/tungstenfs/scratch/gliberal/Users/quingust/Manuscripts/TechnicalPaper/EllenbergLab-MouseEmbryos/Strnad_etal_tracking_data/registered_images_scaled/TestSets/017-*"]


[MultiAggregateOrganoidFeaturesTask]
# datasets to process that are not tracked (lumen/organoid volume only)
movie_dirs=[]


# meshes ###############################################################
[SegmentationMeshTask]

[VolumeGridTask]
raw_channel_subdirs=["mCherry", "EGFP"]
ref_mesh_subdir=cell_mesh
colormaps=["gray", "red"]
blending_mode=max

out_subdir=rgb_grid

[ViewerTask]
movie_dirs=["/tungstenfs/scratch/gliberal/Users/quingust/Manuscripts/TechnicalPaper/EllenbergLab-MouseEmbryos/Strnad_etal_tracking_data/registered_images_scaled/*_*"]
nuclei_seg_subdir=nuclei_segmentation
cell_seg_subdir=cell_segmentation

# tracking #############################################################
[BuildTrackingTrainingRecordTask]
training_base_dir=/tungstenfs/scratch/gliberal/Users/quingust/Manuscripts/TechnicalPaper/EllenbergLab-MouseEmbryos/Strnad_etal_tracking_data/registered_images_scaled/trainingModels/tracking_training/969696_raw/
ch_subdir=mCherry
spacing=(0.52,0.52,0.52)
train_fraction=0.95
valid_fraction=0.05
min_patch_size=(96,96,96,2)
patch_margins=(24,24,24,0)
xml_tree=embryo_mamut.xml


# The task below is useful after the first nuclei segmentation model as been trained, so that new trees can be predicted!
# we should increase the patch size to be able to also see the links when cells divide! (96,96,96,X) is probably too small.
[TrackingTrainingTask]
plot_dataset=true
training_base_dir=/tungstenfs/scratch/gliberal/Users/quingust/Manuscripts/TechnicalPaper/EllenbergLab-MouseEmbryos/Strnad_etal_tracking_data/registered_images_scaled/trainingModels/tracking_training/969696_raw/
movie_dirs=["/tungstenfs/scratch/gliberal/Users/quingust/Manuscripts/TechnicalPaper/EllenbergLab-MouseEmbryos/Strnad_etal_tracking_data/registered_images_scaled/TrainingSets/*-*"
            ]


downsampling_factor=(4,4,4)
n_downsampling_channels=64
n_groups=4
dilation_rates=(1, 2, 4, 8)
channels_per_group=64
n_steps=6
dropout=0.1
n_classes=2
spacing=(0.52,0.52,0.52)

train_batch_size=1
train_batches_per_epoch=200
valid_batch_size=4
epochs=3000
n_restarts=3
learning_rate=0.0001
patch_size=(96,96,96,-1)
suffix=20220511-scratch_raw_3

intra_margin=2.0
inter_margin=6.0
jaccard_hinge=0.3
jaccard_eps=0.1

resume_weights=/tungstenfs/scratch/gliberal/Users/quingust/Manuscripts/TechnicalPaper/EllenbergLab-MouseEmbryos/Strnad_etal_tracking_data/registered_images_scaled/trainingModels/tracking_training/969696_raw/out/RDCNet-F4-DC64-OC10-G4-DR1-2-4-8-GC64-S6-D0.1_20220511-scratch_raw_2/weights_latest.h5

# resume_weights=/tungstenfs/scratch/gliberal/Users/quingust/Manuscripts/TechnicalPaper/EllenbergLab-MouseEmbryos/Strnad_etal_tracking_data/registered_images_scaled/trainingModels/tracking_training/969696_raw/out/RDCNet-F4-DC64-OC10-G4-DR1-2-4-8-GC64-S6-D0.1_20220511-scratch_raw_3/weights_latest.h5

intensity_offset_sigma=0.5
intensity_scaling_bounds=(0.1, 10.)

[NucleiTrackingTask]
ch_subdir=mCherry
out_subdir_nuclei=track_nuclei-20220511-scratch_raw_3
out_subdir_link=track_link-20220511-scratch_raw_3
out_subdir_score=track_score-20220511-scratch_raw_3

[ExtractTrackingFeaturesTask]
out_subdir=track_props-20220511-scratch_raw_3

[BuildTreeTask]
xml_bdv=dataset_raw.xml
max_n_nuclei = 80
min_track_length = 10

[MultiBuildTreeTask]
# movie_dirs=["/tungstenfs/scratch/gliberal/Users/quingust/Manuscripts/TechnicalPaper/EllenbergLab-MouseEmbryos/Strnad_etal_tracking_data/registered_images_scaled/TrainingSets/013-*",
#             "/tungstenfs/scratch/gliberal/Users/quingust/Manuscripts/TechnicalPaper/EllenbergLab-MouseEmbryos/Strnad_etal_tracking_data/registered_images_scaled/TrainingSets/015-*"
#             ]
movie_dirs=["/tungstenfs/scratch/gliberal/Users/quingust/Manuscripts/TechnicalPaper/EllenbergLab-MouseEmbryos/Strnad_etal_tracking_data/registered_images_scaled/TestSets/*-*" 
# "/tungstenfs/scratch/gliberal/Users/quingust/Manuscripts/TechnicalPaper/EllenbergLab-MouseEmbryos/Strnad_etal_tracking_data/registered_images_scaled/TestSets/*-*"
            ]
```
