## Supplementary Software for "Multiscale light-sheet organoid imaging framework": parallel_image_compressor.py

```
""" 
Microscope image compression script for raw tiff images using lzw compression method.

The script performs the following tasks:
- a producer process queues files found within subfolders of <source_path>
    only schedule a total amount of files under 10TiB (default)
- a dispatcher schedules tasks to corresponding consumers
    only schedule a total amount of files under 75GiB (default) in parallel
- a defined number of consumer proceses run in parallel and handling scheduled taks
    check for corrupted files
    copy files to <backup_path>
    compresses TIFF images (using lzw) and copy files back to <source_path>


"""

__author__      = "Dario Vischi"
__copyright__   = "Copyright 2019, " \
                  "Friedrich Miescher Institute for Biomedical Research, " \
                  "Basel, Switzerland"
__version__     = "0.3.1"


from argparse import ArgumentParser
import errno
import hashlib
import logging
import logging.handlers
import multiprocessing as mp
import os
from PIL import Image, ImageSequence
import psutil
import queue
import shutil
import signal
import sys
import time


class Producer(mp.Process):
    def __init__(self, source_path, task_queue, log_queue, total_file_size_limit=None):
        super(Producer, self).__init__()
        self.interrupt_event = mp.Event()
        self.done_event = mp.Event()
        self.source_path = source_path
        self.task_queue = task_queue
        self.log_queue = log_queue

        self.total_file_size_limit = total_file_size_limit

    def stop(self, signal, frame):
        self.interrupt_event.set()

    def run(self):
        self.logger = logging.getLogger('producer_process')
        self.logger.setLevel(logging.DEBUG)
        qh = logging.handlers.QueueHandler(self.log_queue)
        qh.setLevel(logging.DEBUG)
        self.logger.addHandler(qh)

        stop_signals = (signal.SIGINT, signal.SIGTERM, signal.SIGABRT)
        for stop_signal in stop_signals:
            signal.signal(stop_signal, self.stop)

        total_file_size = 0

        self.logger.info("Starting producer process with PID '%s'" % os.getpid())
        for path, subdirs, files in os.walk(self.source_path):
            for file_name in files:
                if file_name.lower().endswith('.tif'):
                    file_path = os.path.join(path, file_name)
                    file_size = os.path.getsize(file_path)

                    try:
                        with Image.open(file_path) as image:
                            file_compression = image.info["compression"]
                            if file_compression == 'tiff_lzw':
                                # to get a copy of the file within the backup path we have to process it
                                pass
                            elif file_compression != 'raw':
                                self.logger.warning(
                                    "Found TIFF file with an unknown compression (%s) '%s'!"
                                    % (file_compression, file_path)
                                )
                                continue
                    except Exception:
                        self.logger.warning(
                            "Found corrupted/invalid TIFF file '%s'!" % file_path
                        )
                        continue

                    total_file_size += file_size
                    self.task_queue.put({
                        'sender': os.getpid(), 'data': {
                            'file_type': 'image',
                            'file_path': file_path,
                            'file_size': file_size
                        }
                    })
                else:
                    file_path = os.path.join(path, file_name)
                    file_size = os.path.getsize(file_path)

                    total_file_size += file_size
                    self.task_queue.put(
                        {
                            'sender': os.getpid(), 'data': {
                                'file_type': 'data',
                                'file_path': file_path,
                                'file_size': file_size
                            }
                        }
                    )

                if (
                    self.total_file_size_limit is not None and
                    total_file_size >= self.total_file_size_limit
                ):
                    self.logger.info(
                        (
                            "Scheduled files with a total size of %s bytes.\n"
                            "Stop collecting further files!"
                        ) % total_file_size
                    )
                    break

                if self.interrupt_event.is_set():
                    break
            if (
                (
                    self.total_file_size_limit is not None and
                    total_file_size >= self.total_file_size_limit
                )or self.interrupt_event.is_set()
            ):
                break

        # handshake with master process
        if not self.interrupt_event.is_set():
            self.task_queue.put(None)
            self.done_event.set()
            self.interrupt_event.wait()

        self.logger.info("Terminating producer process with PID '%s'" % os.getpid())


class Dispatcher(mp.Process):
    def __init__(self, task_queue, notification_queue, log_queue, memory_limit=None):
        super(Dispatcher, self).__init__()
        self.interrupt_event = mp.Event()
        self.done_event = mp.Event()
        self.notification_queue = notification_queue
        self.task_queue = task_queue  # not relyable as soon as producer is dead
        self.log_queue = log_queue

        self.consumer_pid_map = {}
        self.image_consumers = {}
        self.data_consumers = {}
        self.image_tasks = []
        self.data_tasks = []
        self.producer_terminated = False
        self.memory_limit = memory_limit
        self.memory_usage = 0

        self.dispatch_counter = 0

    def subscribe(self, consumer):
        if self.is_alive():
            raise RuntimeError(
                "Consumers can only be subscribed before starting the "
                "dispatcher process!"
            )

        if isinstance(consumer, ImageConsumer):
            self.image_consumers[consumer.pid] = {
                'queue': consumer.task_queue,
                'task': None
            }
            self.consumer_pid_map[consumer.pid] = self.image_consumers[consumer.pid]
        elif isinstance(consumer, DataConsumer):
            self.data_consumers[consumer.pid] = {
                'queue': consumer.task_queue,
                'task': None
            }
            self.consumer_pid_map[consumer.pid] = self.data_consumers[consumer.pid]
        else:
            raise RuntimeError(
                'Try to subscribe consumer of unknown type "%s"!'
                % consumer.__class__.__name__
            )

    @property
    def waiting_image_consumers(self):
        return self.get_consumers(type='image', status='waiting')

    @property
    def running_image_consumers(self):
        return self.get_consumers(type='image', status='running')

    @property
    def waiting_data_consumers(self):
        return self.get_consumers(type='data', status='waiting')

    @property
    def running_data_consumers(self):
        return self.get_consumers(type='data', status='running')

    def get_consumers(self, type=None, status=None):
        consumers = {}
        if type is None:
            consumers.update(self.image_consumers)
            consumers.update(self.data_consumers)
        elif type == "image":
            consumers = self.image_consumers
        elif type == "data":
            consumers = self.data_consumers
        else:
            raise ValueError(
                'Try to find unknown consumer type  "%s"!' % type
            )
        
        if status is None:
            predicate = lambda task: True
        elif status == "waiting":
            predicate = lambda task: task is None
        elif status == "running":
            predicate = lambda task: task is not None
        else:
            raise ValueError(
                'Try to find unknown consumer status  "%s"!' % status
            )

        return {
            pid: properties for pid, properties in consumers.items()
            if predicate(properties['task'])
        }

    def dispatch_notification_messages(self):
        try:
            while True:
                message = self.notification_queue.get(block=False)
                consumer_pid, data = message['sender'], message['data']
                if consumer_pid not in self.consumer_pid_map:
                    raise RuntimeError(
                        'Ignore message "%s" from non-subscribed consumer %s!'
                        % (message, consumer_pid)
                    )
                if data['notification'] == 'job_done':
                    self.logger.debug(
                        'Task successfully completed by consumer "%s"...'
                        % consumer_pid
                    )

                    self.memory_usage -= \
                        self.consumer_pid_map[consumer_pid]['task']['file_size']
                    self.consumer_pid_map[consumer_pid]['task'] = None
                else:
                    raise RuntimeError(
                        'Ignore unexpected message "%s" from consumer %s!'
                        % (message, consumer_pid)
                    )
        except queue.Empty:
            pass

    def dispatch_producer_message(self):
        if self.memory_usage > self.memory_limit:
            self.logger.info(
                "Maximum assigned memory limitation reached. "
                "Throttle distribution..."
            )
            time.sleep(10)
            return

        try:
            if self.dispatch_counter % 100 == 0:
                # clean-up zombie processes
                for consumers, tasks in zip(
                    [self.image_consumers, self.data_consumers],
                    [self.image_tasks, self.data_tasks]
                ):
                    for pid in list(consumers.keys()):
                        consumer = consumers[pid]
                        consumer_process = psutil.Process(pid)
                        if consumer_process.status() == psutil.STATUS_ZOMBIE:
                            if consumer['task'] is not None:
                                # re-schedule un-completed task
                                tasks.put({
                                    'sender': os.getpid(),
                                    'data': consumer['task']
                                })
                            del consumers[pid]
                            del self.consumer_pid_map[pid]

            if not self.producer_terminated:
                message = self.task_queue.get(block=False)
                if message is None:
                    self.producer_terminated = True
                else:
                    producer_pid, data = message['sender'], message['data']
                    if data['file_type'] == 'image':
                        self.image_tasks.append(message)
                    elif data['file_type'] == 'data':
                        self.data_tasks.append(message)
                    else:
                        raise RuntimeError(
                            'Ignore unexpected message "%s" from producer %s!'
                            % (message, producer_pid)
                        )

            for waiting_consumers, tasks in zip(
                [self.waiting_image_consumers, self.waiting_data_consumers],
                [self.image_tasks, self.data_tasks]
            ):
                if len(waiting_consumers) == 0 or len(tasks) == 0:
                    continue

                message = tasks.pop()
                consumer_pid, consumer = next(iter(waiting_consumers.items()))

                producer_pid, data = message['sender'], message['data']

                self.logger.debug(
                    'Dispatching file "%s" to consumer "%s"...'
                    % (data['file_path'], consumer_pid)
                )

                consumer['task'] = data
                consumer['queue'].put({'sender': os.getpid(), 'data': data})

                # for simplicity, we exaggerate the memory usage for data tasks
                # copying a file using shutil requires only 16384 bytes - not the full file size!
                # https://blogs.blumetech.com/blumetechs-tech-blog/2011/05/faster-python-file-copy.html
                self.memory_usage += data['file_size']
        except queue.Empty:
            pass

    def stop(self, signal, frame):
        self.interrupt_event.set()

    def run(self):
        self.logger = logging.getLogger('dispatcher')
        self.logger.setLevel(logging.DEBUG)
        qh = logging.handlers.QueueHandler(self.log_queue)
        qh.setLevel(logging.DEBUG)
        self.logger.addHandler(qh)

        stop_signals = (signal.SIGINT, signal.SIGTERM, signal.SIGABRT)
        for stop_signal in stop_signals:
            signal.signal(stop_signal, self.stop)

        self.logger.info("Starting dispatcher with PID '%s'" % os.getpid())
        while not self.interrupt_event.is_set():
            # health check
            for consumers, consumer_type in zip(
                [self.image_consumers, self.data_consumers] , ['image', 'data']
            ):
                if len(consumers) == 0:
                    self.logger.critical(
                        "No (alive) %s consumer available for dispatching!"
                        % consumer_type
                    )
                    self.interrupt_event.is_set()
                    continue

            self.dispatch_counter += 1
            try:
                self.dispatch_notification_messages()
                self.dispatch_producer_message()
                #time.sleep(1)
            except RuntimeError as ex:
                self.logger.warning(str(ex))

            if (
                self.producer_terminated and
                len(self.image_tasks) + len(self.data_tasks) == 0 and
                len(self.running_image_consumers) + len(self.running_data_consumers) == 0
            ):
                break

        # handshake with master process
        if not self.interrupt_event.is_set():
            for pid, consumer in self.consumer_pid_map.items():
                consumer['queue'].put(None)
            self.done_event.set()
            self.interrupt_event.wait()

        self.logger.info("Terminating dispatcher with PID '%s'" % os.getpid())


class ImageConsumer(mp.Process):
    def __init__(self, source_path, backup_path, notification_queue, log_queue):
        super(ImageConsumer, self).__init__()
        self.interrupt_event = mp.Event()
        self.source_path = source_path
        self.backup_path = backup_path
        self.task_queue = mp.Queue()
        self.notification_queue = notification_queue
        self.log_queue = log_queue

    def stop(self, signal, frame):
        self.interrupt_event.set()

    def run(self):
        logger = logging.getLogger('image_consumer_process')
        logger.setLevel(logging.DEBUG)
        qh = logging.handlers.QueueHandler(self.log_queue)
        qh.setLevel(logging.DEBUG)
        logger.addHandler(qh)

        stop_signals = (signal.SIGINT, signal.SIGTERM, signal.SIGABRT)
        for stop_signal in stop_signals:
            signal.signal(stop_signal, self.stop)

        logger.info("Starting image consumer process with PID '%s'" % os.getpid())
        while not self.interrupt_event.is_set():
            try:
                message = self.task_queue.get(block=True, timeout=1)
                if message is None:
                    self.interrupt_event.set()
                    continue
            except queue.Empty:
                continue

            sender, data = message['sender'], message['data']
            image_path = data['file_path']

            logger.debug('Processing image "%s"...' % image_path)

            image_backup_path = os.path.join(
                self.backup_path,
                image_path[len(self.source_path)+1:]
            )
            try:
                os.makedirs(os.path.dirname(image_backup_path))
            except (OSError, FileExistsError) as err:
                # directory may already exists...
                if err.errno != errno.EEXIST:
                    raise

            try:
                shutil.copy2(image_path, image_backup_path)
                # shutil.copy(image_path, image_backup_path)    # copy less metadata
                
                image = Image.open(image_backup_path)  # check for corrupted files

                # do compression...
                image_pages = []
                for page in ImageSequence.Iterator(image):
                    image_pages.append(page.copy())

                image_pages[0].save(
                    image_path, compression="tiff_lzw", save_all=True,
                    append_images=image_pages[1:]
                )

                for page in image_pages:
                    page.close()
                del image_pages[:]
                image.close()

                with Image.open(image_path) as tmp_image:
                    # check for corrupted files
                    pass

            except Exception as ex:
                logger.warning(
                    'An unexpected error occurred while compressing '
                    'image "%s":\n%s' % (image_path, str(ex))
                )

            self.notification_queue.put({
                'sender': os.getpid(),
                'data': {'notification': 'job_done'}
            })

        logger.info("Terminating image consumer process with PID '%s'" % os.getpid())


class DataConsumer(mp.Process):
    def __init__(self, source_path, backup_path, notification_queue, log_queue):
        super(DataConsumer, self).__init__()
        self.interrupt_event = mp.Event()
        self.source_path = source_path
        self.backup_path = backup_path
        self.task_queue = mp.Queue()
        self.notification_queue = notification_queue
        self.log_queue = log_queue

    def stop(self, signal, frame):
        self.interrupt_event.set()

    def run(self):
        logger = logging.getLogger('data_consumer_process')
        logger.setLevel(logging.DEBUG)
        qh = logging.handlers.QueueHandler(self.log_queue)
        qh.setLevel(logging.DEBUG)
        logger.addHandler(qh)

        stop_signals = (signal.SIGINT, signal.SIGTERM, signal.SIGABRT)
        for stop_signal in stop_signals:
            signal.signal(stop_signal, self.stop)

        logger.info("Starting data consumer process with PID '%s'" % os.getpid())
        while not self.interrupt_event.is_set():
            try:
                message = self.task_queue.get(block=True, timeout=1)
                if message is None:
                    self.interrupt_event.set()
                    continue
            except queue.Empty:
                continue

            sender, data = message['sender'], message['data']
            file_path = data['file_path']

            logger.debug('Processing file "%s"...' % file_path)

            file_backup_path = os.path.join(
                self.backup_path,
                file_path[len(self.source_path)+1:]
            )
            try:
                os.makedirs(os.path.dirname(file_backup_path))
            except (OSError, FileExistsError) as err:
                # directory may already exists...
                if err.errno != errno.EEXIST:
                    raise

            try:
                shutil.copy2(file_path, file_backup_path)
            except Exception as ex:
                logger.warning(
                    'An unexpected error occurred while copying '
                    'file "%s":\n%s' % (file_path, str(ex))
                )

            # Check file integrity
            source_md5 = hashlib.md5()
            with open(file_path, 'rb') as source_fh:
                for buffer in iter(lambda: source_fh.read(4096), b''):
                    source_md5.update(buffer)

            target_md5 = hashlib.md5()
            with open(file_backup_path, 'rb') as target_fh:
                for buffer in iter(lambda: target_fh.read(4096), b''):
                    target_md5.update(buffer)

            if source_md5.hexdigest() != target_md5.hexdigest():
                logger.warning(
                    'Found corrupted backup of file "%s"!' % (file_path,)
                )

            self.notification_queue.put({
                'sender': os.getpid(),
                'data': {'notification': 'job_done'}
            })

        logger.info("Terminating data consumer process with PID '%s'" % os.getpid())


class Logger(mp.Process):
    def __init__(self, log_queue):
        super(Logger, self).__init__()
        self.interrupt_event = mp.Event()
        self.log_queue = log_queue

    def stop(self, signal, frame):
        self.interrupt_event.set()

    def run(self):
        logger = logging.getLogger('logger_process')
        logger.propagate = False
        logger.setLevel(logging.INFO)

        formatstr = (
            '%(asctime)s %(name)s[%(process)s] %(levelname)s: %(message)s'
        )
        formatter = logging.Formatter(formatstr)

        sh = logging.StreamHandler(sys.stdout)
        sh.setLevel(logging.INFO)
        sh.setFormatter(formatter)
        logger.addHandler(sh)

        stop_signals = (signal.SIGINT, signal.SIGTERM, signal.SIGABRT)
        for stop_signal in stop_signals:
            signal.signal(stop_signal, self.stop)

        logger.info("Starting logger process with PID '%s'" % os.getpid())
        while not self.interrupt_event.is_set():
            try:
                record = self.log_queue.get(block=False, timeout=0)
            except queue.Empty:
                time.sleep(1)
                continue

            logger.handle(record)


if __name__ == '__main__':
    """
    if os.name != 'posix':
        raise RuntimeError(
            "Image Compressor only runs on POSIX compatible operating systems!"
        )
    """

    parser = ArgumentParser()
    parser.add_argument(
        'source_path',
        help='source path to process.'
    )
    parser.add_argument(
        'backup_path',
        help='backup path where to move raw data.'
    )
    parser.add_argument(
        '--processes',
        type=int,
        default=mp.cpu_count(),
        help='number of parallel consumer processes.'
    )

    args = parser.parse_args()
    source_path = (
        args.source_path if not args.source_path.endswith(os.sep)
        else args.source_path[:-1]
    )
    backup_path = (
        args.backup_path if not args.backup_path.endswith(os.sep)
        else args.backup_path[:-1]
    )
    processes = args.processes

    task_queue = mp.Queue()
    notification_queue = mp.Queue()
    log_queue = mp.Queue()

    logger = logging.getLogger('main_process')
    logger.setLevel(logging.INFO)
    qh = logging.handlers.QueueHandler(log_queue)
    qh.setLevel(logging.INFO)
    logger.addHandler(qh)
    
    logger_process = Logger(log_queue)
    producer_process = Producer(
        source_path, task_queue, log_queue,
        total_file_size_limit=16492674416639  # = 15TiB
    )
    dispatcher_process = Dispatcher(
        task_queue, notification_queue, log_queue,
        memory_limit=187904819200  # = 175 GiB
    )
    image_consumer_processes = [
        ImageConsumer(source_path, backup_path, notification_queue, log_queue)
        for _ in range(processes)
    ]
    data_consumer_processes = [
        DataConsumer(source_path, backup_path, notification_queue, log_queue)
        for _ in range(2)
    ]
    consumer_processes = image_consumer_processes + data_consumer_processes
    processes = [logger_process, producer_process, dispatcher_process] + consumer_processes

    for process in processes:
        process.daemon = True  # has no effect as all sub-processes ignore the interrupt signal

    def interrupt_handler(*args):
        print('Terminating process after a keyboard interrupt...')
        exit(1)

    stop_signals = (signal.SIGINT, signal.SIGTERM, signal.SIGABRT)
    for stop_signal in stop_signals:
        signal.signal(stop_signal, interrupt_handler)

    logger_process.start()
    time.sleep(1)
    for consumer_process in consumer_processes:
        consumer_process.start()
        dispatcher_process.subscribe(consumer_process)
    producer_process.start()
    dispatcher_process.start()
    time.sleep(1)

    # monitoring processes
    while producer_process is not None or dispatcher_process is not None:
        if producer_process is not None:
            if producer_process.is_alive() and producer_process.done_event.is_set():
                producer_process.interrupt_event.set()  # free process
                producer_process = None
            elif not producer_process.is_alive():
                logger.critical("Producer process was killed!")
                os.kill(os.getpid(), signal.SIGINT)
        if dispatcher_process is not None:
            if dispatcher_process.is_alive() and dispatcher_process.done_event.is_set():
                dispatcher_process.interrupt_event.set()  # free process
                dispatcher_process = None
            elif not dispatcher_process.is_alive():
                logger.critical("Dispatcher process was killed!")
                os.kill(os.getpid(), signal.SIGINT)
        if not logger_process.is_alive():
            raise RuntimeError("Logger process was killed!")

        time.sleep(1)

    for consumer_process in consumer_processes:
        consumer_process.interrupt_event.set()
        consumer_process.join()

    time.sleep(1)

    logger_process.interrupt_event.set()
    logger_process.join()
```
