## Supplementary Software for "Multiscale light-sheet organoid imaging framework": README.txt

Supplementary Code Information for "Multiscale light-sheet organoid imaging framework" from de Medeiros, Gustavo, Ortiz, Raphael, Strnad, Petr, Boni, Andrea, Moos, Franziska, Repina, Nicole, Chalet-Meylan, Ludivine, Maurer, Francisca, & Liberali, Prisca. 

This file contains:


1) elephant_mouse_models: contains the linking and spot models trained for tracking prediction for mouse data from Strnad et al. Nature Methods (2015) using the Elephant Tracker

2) lstree_mouse_models: contains the lstree based nuclei segmentation and tracking prediction models for mouse data from Strnad et al. Nature Methods (2015)

3) Elastix_parameters_Affine.txt: parameters file to be used with elastix for registration of fixed organoids to their last position during live imaging

4) parallel_image_compressor.py: python script for compressing images in parallel using the lzw lossless standard

5) rescale_uint8-uint16.ipynb: jupyter notebook for rescaling 8bit into 16bit images during the pre-processing for the mouse data from Strnad et al. Nature Methods (2015)

6) Stardist_config.json: configuration file containing parameters used for nuclei segmentation prediction using Stardist
