## Supplementary Software for "Multiscale light-sheet organoid imaging framework": rescale_uint8-uint16.ipynb

```
# %% [markdown]
## Rescaling images from 8bit to 16bit
this small notebook, via single core use, rescales uint8 images into uint16 using tifffile library.

# %% [code]
from tifffile import imread, imwrite
import numpy as np
import os

# %% [code]
base_path = '/tungstenfs/scratch/gliberal/Users/quingust/Manuscripts/TechnicalPaper/EllenbergLab-MouseEmbryos/Strnad_etal_tracking_data/registered_images_scaled/old/TEST'
path_uint8 = os.path.join(base_path, 'uint8_images' )     # considering images in uint8_images folder
path_uint16 = os.path.join(base_path, 'uint16_images' )   # this folder will be created in case it does not exist


files_uint8 = os.listdir(path_uint8)
print(files_uint8)

for x in files_uint8:
    
    print(x)
    img8 = imread(os.path.join(path_uint8, x))
    img16 = img8.astype(np.uint16)
    imwrite(os.path.join(path_uint16,x),img16)

# %% [code]
```
