## Supplementary Information for "Multiscale light-sheet organoid imaging framework"

^2^Current address: Disney Research Studios, Stampfenbachstrasse 48, 8006 Zürich

^3^Current address: Viventis Microscopy Sàrl, EPFL Innovation Park, Building C, 1015 Lausanne.

### Summary Note

Here we provide technical information about the rationale behind the imaging steps (**Supplementary Note 1**), pre-processing (**Supplementary Note 2**) lineage tree and segmentation (**Supplementary Note 3**) and feature extraction and data visualization (**Supplementary Note 4**) pipelines inside the LSTree framework (https://github.com/fmi-basel/LSTree). **Supplementary Note 5** discusses the validation strategies for our nuclei and tree prediction approaches. And highlights the challenges and care needed for using LSTree on other imaging data than that from the organoids presented in the main text (using mouse data from a 2015 publication). In **Supplementary Note 6** we discuss the challenges regarding fixation and backtracking. **Supplementary Table 1** shows details on the features extracted for **Figure 1f**, whereas **Supplementary Table 2** provides an overall list of extracted features with the feature extraction task from LSTree. **Supplementary Figures** and **Supplementary Movies** captions can be found at the end.

Please not that all code used in this work can be found in the **Supplementary Code** and in the **LSTree repository** (<https://github.com/fmi-basel/LSTree>). For any issues, questions or comments, please use the issues section from LSTree repository (https://github.com/fmi-basel/LSTree/issues) or contact Gustavo de Medeiros directly

### Supplementary Note 1: Imaging steps

For all live imaging experiments, we have performed routine steps in order to improve number of organoids imaged as well as general image quality of the recorded samples.

**Position dependent illumination alignment**

We place the cell containing Matrigel drops in FEP foil compartments created with a stamp (Supplementary Figure 1), each compartment being 1.5X1.5 mm^2^ and 1 mm deep. After placing the sample holder in the microscope, a first initial step is to try to find as many positions as possible by searching for healthy cells using transmitted light. In some cases also short exposure fluorescence images can be used to see whether fluorescent signal is strong enough (we kept a value of close to 1000 gray values as a good intensity measure, considering the camera background to be around 110). Among the visual criteria for accepting a cell as a new position for imaging were nuclear integrity, shape, and intensity. After all initial positions were defined, a position dependent illumination alignment was performed. For this, we looped over all previously defined positions once more and for each one set the best offset values for each illumination arm so that both illumination planes would be at best coplanar with the imaging plane, as well as improving general image contrast. Currently this process is performed manually, and for 30 positions the necessary time from placing the sample holder onto the microscope to pressing the “Run” button to start the timelapse is around 1.5 hours. Once aligned, each particular position would also have extra values for offsetting the stages and galvanometric mirrors in order to provide the initially desired offset, and he recording can start.

Generally speaking, the maximum number of positions possible to record depends on the exposure time, stack size and time interval between time-points. Furthermore, since medium change happens while acquisition is running, during the interval between time-points, it is wise to leave around 45 seconds to 1 minute of time for pipetting. Taken together, this means that in general around 30 positions can be imaged per experiment.

Acquired images are compressed on the fly in a lossless manner using LZW compression. Since LZW compression is relatively fast, our implementation allows data compression as the microscope runs, so that neither storage overhaul nor post-acquisition processing is needed. The compression ratio is content dependent and typically higher on images with more background pixels and higher signal-to-noise ratio (SNR). Overall, we achieve a compression ratio of at least 1/3 for the uncropped stacks. Note: although very useful ultimately for storage, compression is not a pre-requisite for any of the processing steps later.

**Important:** Many factors change the image quality during acquisition through many days; even medium change can have an effect on the amount of diffusive light detected by the microscope (as exemplified in **Supplementary Figure 2**). Furthermore, the objective configuration of the light-sheet microscope, although allowing for direct access to samples especially close to the bottom of the sample holder, also has its disadvantages: due to a single detection objective, typically organoids with diameter larger than 100 μm (Day 4/5 from single cells) will not be imaged across with the same image quality, and cells on the far side of the sample become consistently more blurred as it further grows. Even with tiling, a thorough analysis via tracking and segmentation becomes virtually impossible, as also demonstrated with large cerebral organoids in He, Maynard, Jain, Gerber et al. *Nature Methods* (2022).

### Supplementary Note 2: Pre-processing pipeline

Here we present the main contributions for achieving high-quality yet low storage stacks of live organoid recordings.

**Cropping**

The initial field of view (FOV) of the camera is kept relatively large to avoid losing samples due to drift during long-term acquisition. Late-stage organoids occupy about ¼ of the acquired area (2048X2048 pixels) and are typically not centered over the entire movie. To reduce memory footprint and reduce downstream processing time, the organoids are tracked and cropped in a combined step (**Figure 2a**, **Supplementary Figure 3**).

Organoids' bounding boxes are first determined on the nuclei channel and independently for each frame using x,y and z maximum intensity projections (MIPs). Since multiple organoids might appear in the field of view (especially at early time-points), the largest object (or a manually selected object) on the last frame is tracked backward in time by finding its closest match in the previous frame until the first frame is reached. The minimum crop size required for the entire movie is then computed along each axis. At this point crops are reviewed with our interactive tool showing orthogonal MIPs with overlaid bounding boxes of all detected objects and of the current global bounding box. Manual corrections are made for instance to account for stage movements during medium change. After each correction, the linking process is updated and overall, all manual operations can be done within a few minutes. Finally, all time-points and channels are cropped by centering the global bounding box on the tracked organoid. These cropped stacks are then used in all subsequent processing and original full-FOV images can be archived on tape or discarded to reduce storage requirement.

**Denoising and deconvolution**

Live imaging oftentimes requires low-light signal resulting in low SNR which is problematic for traditional deconvolution algorithms. We therefore first remove pixel-wise independent noise with a fully convolutional neural network (FCN) trained with the Noise2Void scheme^1^ on a few randomly selected frames from each movies/channel. Each 2D slice of a 3D stack is processed independently and Fourier frequencies with amplitude higher than the input are clipped. After denoising, background levels are corrected by subtracting the minimum intensity projection along z, assuming that for each pixel the background is visible on at least one of the z-slice. Finally, image intensities are rescaled based on min/max bounds over the whole movie to make use of the entire 16-bit integer format in which the outputs are saved.

Denoised and background-corrected images are then deconvolved with a measured point spread function (PSF) using a TensorFlow implementation of Richardson-Lucy algorithm (<https://github.com/hammerlab/flowdec>). Throughout this work we used measured and averaged PSFs using Huygens PSF-Distill from 0.25 µm beads (ThermoFisher, Cat-No T7280) acquired with our light-sheet microscope.

As mentioned in the main text, the segmentation strategies proposed in this work do not necessarily require denoised and deconvolved data as input, these are rather optional steps for training of the models, which may aid in providing better segmentation outputs. In practice, denoising and deconvolution of the raw images proved itself especially useful during the correction of predicted trees using Mastodon. For a more thorough discussion on utilizing LSTree for different datasets / image types, please refer to **Supplementary Note 5,** where we evaluate the validation of LStree with different prediction approaches / different datasets.

### Supplementary Note 3: Lineage tree and segmentation prediction pipelines

**Overall strategy**

Nuclei tracking seeds serve as the foundation to obtain temporally coherent full nuclei/cells segmentation masks. To leverage this more easily obtained sparse information, deep learning models are trained to fully segment the nuclei/cells with a limited number of additional complete annotations. Furthermore, once the initial data have been processed, the outputs are fed back in a loop to train a joint tracking/segmentation model that greatly reduces the time required to produce curated tracking seeds.

**Initial lineage tracking / manual curation of predicted trees**

Initial lineage trees are obtained by importing the deconvolved image for manual tracking in Mastodon FIJI Plugin (<https://github.com/mastodon-sc/mastodon>). One main strength of Mastodon is that it provides tools for automatic tracking. However, for long time-lapse imaging the quality of the images usually change over time, as well as the density of nuclei, their relative movement, etc. As a result, these changes enforce the need to change the initial parameters set for the automatic tracking, as otherwise the automated tracker either jumps from nucleus A to nucleus B, or even simply stops. On extreme cases, the changes of the available parameters of Mastodon may still not suffice on reliable automated tracking, and so manual curation / tracking becomes necessary. This is especially the case for later timepoints, as exemplified in the case for tracking the fate of the progeny of the initially merged nuclei presented in **Figure 5f.**

After an initial round, lineage trees are automatically predicted and exported as a .xml file compatible with Mastodon. Additionally, plots of the predicted trees with overlaid nuclei volume provide a guide to focus the manual curation on potential mistakes. Other metrics include linking distance between two seeds and the level of overlap to parent from the segmented nuclei (**Supplementary Figure 4**).

Alternatively, for the cases where not enough imaging data is present to generate a robust tree prediction model via RDCNet, it is also possible to use Elephant (<https://elephant-track.github.io/#/v0.3/>) as a means to get the lineage tree output file as MaMuT.xml and continue with the segmentation steps via LSTree.

**Deep learning model training**

All deep learning models are trained in TensorFlow on a Nvidia Quadro RTX 6000 GPU with 24 GB VRAM using the Adam optimizer and cosine scheduler with learning rates from 10^−4^

to 10^−6^. The complete list of training hyperparameters used for each model/task (architecture definition, patch size, batch size, data augmentation, train/validation split, etc.) can be found in the main config file of the source code, and a brief explanation for each of these parts is present in the Github repository.

**Nuclei segmentation**

Nuclei are segmented in 3D by expanding on a previously reported method: RDCNet^2^ (https://github.com/fmi-basel/RDCNet). The fully convolutional network (FCN) is trained with a mix of 52 fully annotated and 4500 partially annotated stacks with spheres drawn at the center of each tracked nuclei. During training, the datasets are balanced by repeating the full annotations in random order such that each mini-batch is supervised by an equal number of fully/partially annotated images. Once the model is trained, nuclei segmentations are obtained by assigning the predicted foreground pixel embeddings to their closest tracking seeds’ embedding which enforces the correct number of nuclei and hence temporal consistency.

Note that in practice one can start training with fewer annotations and go over a few correction-retraining cycles. This is more effective since depending on the image input more or less annotated stacks may need to be annotated and the annotators’ time is focused on most challenging samples (odd-shaped nuclei, poorer image quality, etc.).

**Cell and lumen segmentation**

Cell and lumen segmentation also expand on the RDCNet method. The semantic branch predicts 3 classes, background, lumen, epithelium and is supervised by manual annotations of 86 frames covering all datasets. No manual annotations of individual cells are required. Instead, the previously segmented nuclei are used as partial annotations under the assumption that they are randomly distributed within the cell compartment. Labels of nuclei belonging to multi-nucleated cells are merged based on the tracking information. Since only the membrane channel is provided as input, the network is forced to learn to segment cells. In practice nuclei are not completely randomly distributed (e.g. corners, tapered elongated cells). We therefore also add a regularization term that encourages voxels over the epithelium mask to be assigned to one of the cells (without enforcing which one).

**Joint nuclei segmentation and tracking**

Theoretically, the goal to achieve lineage tracing would be to train a model end-to-end from the complete series of raw images and capture the global context of a tree. It is however not feasible due to memory and computational constraints. While the tracking by detection paradigm allows linking sparse seeds with some global optimization, it becomes impossible to recover from the temporarily inconsistent detection (e.g. missing/extra nuclei) on densely packed cells. Instead, as described in the main text, nuclei are tracked by predicting their segmentation on 2 consecutive frames at the same time, whereby the same nucleus over the time axis is considered as a single instance, i.e. nuclei in the second frame should have the same label as their parent, including nuclei that just divided.

A RDCNet model having 2 channels as input (pseudo 4D) is applied to 2 consecutive frames from the H2B channel. After the last 3D convolutional layer, the output is reshaped as a true 4-dimensional output and the embedding loss is applied normally, with the instances being a nucleus at time *t* plus its daughter(s) at *t+1*. The model is supervised by pairs of frames of the previously predicted nuclei segmentation where the labels of the second frame are changed to match the parent nuclei using the tracking information. During inference all frames are processed by applying the model with a 2-frame wide moving window. The tree is built by linking nuclei across timepoints based on the segmentation overlap as depicted in **Figure 2c**. As this approach produces acyclic graphs, the final edge connecting merged tracks is added by hand in case of a merging event.

In order to quantify the lineage tracking capability, 2 of 7 tracked movies are removed from the training set and a model is trained from scratch with the remaining ones. The predicted trees are then mapped onto the reference ones (i.e. manually corrected ones) to measure the precision, recall and f-1 score of predicted nodes and edges. Each node of the predicted tree is mapped to its closest nuclei in the reference tree. For any given node in the reference tree, the first match is counted as true positive and extra matches as false positives. Late timepoints that cannot be fully tracked, even by hand due the lower image quality are excluded from this analysis.

|  | **DS1: 350 time-points,**  **53 nuclei** | **DS6: 350 time-points,**  **231 nuclei** |
| --- | --- | --- |
| **node precision** | 0.997 | 0.994 |
| **node recall** | 0.998 | 0.996 |
| **node f-1 score** | 0.998 | 0.995 |
| **edge precision** | 0.994 | 0.990 |
| **edge recall** | 0.994 | 0.991 |
| **edge f-1 score** | 0.994 | 0.990 |

Supplementary Table 1: Evaluation of the tree prediction for dataset 006.

With f-1 scores >=0.995 and >0.990 for nodes (i.e. nuclei) and edges respectively, and including many correctly predicted cell divisions, large portion of the tree can be correctly reconstructed and visually identifying topographical mistakes becomes immediate. In addition, overlaying the nuclei volume also highlights positions that require manual corrections. Ultimately time-points are reached where the predicted nuclei masks become less consistent, i.e. they depend more on the joint neighboring frame as evident from the decrease overlap to parent (**Supplementary Figure 4**). This is due to the higher cell density and worse image quality (low SNR, shadowing artifacts) at later time-points. The time of this transition can vary greatly as illustrated from these 2 datasets and ranges from a 40-cell organoid to >200 cells. While there is still an intermediate regime where hand-tracking is more effective on challenging images, our automated lineage tracing method greatly speeds-up the process in general and we expect this gap to be closed as more training data becomes available. However, it is important to note here that, as with any neural network, best results will be achieved if the network is trained on a very specific time-window of organoid development. In our approach we trained models based on lineage trees that initially spanned the first 3 days of growth, and these models are suitable for later retraining to also be able to best encompass days 4 and 5.

**Loss function calculations**

Formally, the loss function to supervise the training of nuclei, cell and tracking follows the same hinged version of the embedding loss function described in [Ortiz et al, MICCAI-MLMI (2020)] where embeddings $y$ are converted into a heatmap for pixel $u$ being part of instance $k$ as:

$$P\left( u=k \right)=\left\{ \begin{aligned} \left[ 1-\frac{\left\| y_{u}-\hat{y}_{k} \right\|-\delta_{intra}}{\delta_{inter}-\delta_{intra}} \right]_{-}, &u\in S_{k} \\ \left[ 1-\frac{\left\| y_{u}-\hat{y}_{k} \right\|-\delta_{intra}}{\delta_{inter}-\delta_{intra}} \right]_{+}, &u\notin S_{k} \end{aligned} \right.$$

Supplementary Equation 1: Embedding loss function used for all training evaluations.

where $\left[ x \right]_{+}=max(0,x)$ and $\left[ x \right]_{-}=min(1,x)$ are the hinges, $\delta_{intra}$ the maximum margin between embeddings of the same instance, $\delta_{inter}$ is the minimum margin between embeddings of different instances and the centroids $\hat{y}_{k}$ are estimated as the mean embedding under the $true$ mask of instance $k$ denoted with $S_{k}$. This generates a heatmap for each instance that can be compared to the one-hot encoded $true$ segmentation. Note that heatmaps are bounded to $\left[ 0,1 \right]$ only where the correct instances are predicted which mitigates vanishing gradient problems but can still be optimized with the soft Jaccard loss without additional modifications.

Pixels outside of the annotated instance masks are supervised by minimizing a new regularization term:

$$R\left( u \right)=1-\left[ \max_{k \in[1,\ldots,n]} P\left( u=k \right) \right]_{-}$$

Supplementary Equation 2: Regularization function for supervising annotated instance masks.

The above minimization pushes all embeddings in unsupervised regions close to one of the instances. In the case of tracking the main difference is that an instance is a nuclei plus its daughter(s) from the next frame so that proper linking can be generated.

### Supplementary Note 4: feature extraction and data visualization

**Organoid level features: medium dependent organoid and lumen swelling**

Organoid-level features such as epithelium, lumen volumes are measured for all datasets. We observed abrupt increase both in lumen and organoid volumes at certain moments of the recordings and generally spaced around 24 hours apart. Concomitant to this, a closer look at the data also shows an increase in signal to noise as well as a general swelling of the tissue, as if part of the autofluorescence from the lumen had been “washed out”. In fact, these moments correspond to the timings of medium exchange, providing the organoids with fresh medium so they can further grow (**Supplementary Figure 3**). Although recent work has also shown expansion, subsequent rupture and contraction of organoids throughout time^3^, the dynamics observed here seem less abrupt and so we hypothesize that this can be just due to the sudden osmotic change between lumen and medium contents, being later equilibrated without organoid rupture~~.~~

**Cell-level features**

Cell-level features are extracted for all tracked datasets. General graph-based features such as generation and displacement are readily available from the tree. Many other features rely on the segmented data. For example, with the nuclei and cells segmentation at hand we can also measure their volume ratio, as exemplified in **Figure 1e**. As illustrated in the **Figure 3c** the nuclei distance to the basal and lumen membranes are measured as the mean distance of nucleus pixels to their closest point on the basal and lumen boundaries respectively. Furthermore, to investigate cell inter-mixing (presented e.g. in **Figure 4g**), we extract a list of neighbors for each cell. This is done by downsampling the image, making the segmentation boundaries of a certain cell to overlap with the boundaries of the closest neighbors. Cells are considered neighbors if their segmentation boundaries are 2 µm apart or less. This corresponds to 1 slice, i.e. maximum sampling in z, and can thus be improved if data is acquired at higher sampling values.

**Digital Organoid Viewer**

To evaluate the evolution of spatial features over time, we developed a web-based viewer having a synchronized view of the tree (with overlaid color) linked to a 3D rendering of the organoid. It also includes orthogonal views of the deconvolved images to rapidly check the fidelity of the cells and nuclei segmentations. To be responsive, the viewer relies on smooth 3D meshes pre-generated from the segmented stacks with VTK (<https://vtk.org/>). The viewer itself is implemented using the python library Holoviews (<https://holoviews.org/>) for the interactive plot of the tree and vtk.js (<https://kitware.github.io/vtk-js>) for the 3D rendering. The viewer can be neatly accessed as stand-alone application from a web browser but also as a jupyter notebook, allowing interactive exploratory analysis.

There is an example usage of the webvier in the LSTree GitHub repository, which uses the viewer to visualize two example datasets using an example notebook.

### Supplementary Note 5: Nuclei segmentation and Tracking prediction validation

**Nuclei Segmentation**

All segmentation strategies used in LSTree are modifications based in the previously published RDCNet work. To validate the nuclei segmentation performance considering all light-sheet training datasets used in this work, we compared our nuclei segmentation with Stardist (<https://github.com/stardist/stardist>). Training datasets comprised of the same 52 fully annotated datasets used for LSTree model training, and we added 10 more fully hand-annotated stacks (randomly selected, yet allowing a good distribution among the different timepoints) for being used as a gold standard.

Following the documentation of Stardist, we first trained a model with the parameters as present in the Stardist ‘Stardist_config.json’ file in the Supplementary Source Code. We then used the trained network to predict the 10 hand-annotated stacks and used several validation metrics (precision, recall, accuracy and f1) to compare its output with our modified RDCNet approach (**Supplementary Figure 4a,b**). Important to note that the comparison between RDCNet and Stardist needs to be taken with care, as currently our RDCNet implementation uses more augmentation steps (especially considering intensity variations) then the Stardist approach (90 rotations around Z only). Thus, the comparison presented here only represents the output one gets by training each standard network without further compromise.

**Tracking Prediction**

For the tracking validation, we utilized LSTree from scratch on mouse embryo data acquired with a different light-sheet microscope and published in 2015 ^4^ (kindly provided by the authors). This data is very different from our organoid data, comprised of isotropic stacks and so we ran the entire workflow from scratch. Because for these datasets we did not possess the PSF of the system, we decided to run the nuclei segmentation training task with the raw data directly, skipping the pre-processing with denoising and deconvolution (for more information please refer to https://github.com/fmi-basel/LSTree/tree/main/example). Since the data provided to us were 8bit images, we had to rescale them to 16bit before usage (a simple jupyter notebook which can perform this task is available in the source code). Furthermore, considering that the previous publication also had lineage trees for all of the datasets, we modified the tracking output to match MaMuT .xml format using a KNIME workflow (<https://kni.me/w/mfKibFKmLuYWuDz4>). Then we hand-annotated 6 randomly selected stacks from 6 different datasets to have dense annotations, and used the spot information from all 6 lineage trees to create sparse annotations for training of a nuclear segmentation prediction model. After this initial training step we trained a tree-prediction model based on all 6 lineage trees and the new nuclei segmentation model, and tested it on 4 datasets.

On this particular case, training of nuclei segmentation turned out to be more involving than for the organoid images, as the already background subtracted movies show very low signal t noise. Therefore, as expected, we had to manually hand-annotate more stacks than for the organoid scenario (63 stacks for the mouse in comparison to 52 previously for the organoid data) and trained nuclei segmentation and tracking prediction models, which are present in the Supplementary Code. As a first result, we managed to get nuclei segmentations which were tendentially smaller than the nuclei themselves (**Supplementary Figure 4c**), and we believe that further training of the network with more hand-annotated data would be beneficial for a better segmentation result.

Nonetheless, we moved on and trained a tracking prediction model based on the previous nuclei segmentation results. As can be seen in **Supplementary Figure 4**, we also further compared the output of tree prediction to that of a trained models for Elephant Tracker (elephant link here). To this end we first trained a Spot prediction model starting from the intensity based “Default model”, which was taking five days of prediction and correction of the model. The Spot prediction model was trained on a dataset that was not later used to compare LSTree with Elephant. For tracking the cells we used the pre-trained “Versatile”-linking model with optical flow support (both Elephant models can be found in **Supplementary Code**). Comparing with the ground truth tracks from the previous publication, both LSTree and Elephant prediction yield rather good results, with Elephant showing less missing values (10 VS 297 for LSTree, **Supplementary Figure 4e**). This is expected due to the fact that the initial segmentation is not yet performing well especially at later timepoints. Furthermore, as the output from the spot prediction of Elephant comprises of not only spheres but ellipsoids, they already offer more spatial information than the expanded spheres used as weak annotations in LSTree, and we assume that this, especially in the case of low SNR imaging, may aid in the tracking prediction, as the ellipsoids will have more chance of overlap than suboptimal segmentations. It is important to mention here that in the case where the tree prediction from LSTree is not good enough, the output tree from Elephant can also be used. Below are the validation values for Elephant and LSTree on the dtaset present in **Supplementary Figure 4.**

|  | **LSTree** | **Elephant** |
| --- | --- | --- |
| **node precision** | 0.969 | 0.939 |
| **node recall** | 0.972 | 0.999 |
| **node f-1 score** | 0.970 | 0.968 |
| **edge precision** | 0.969 | 0.938 |
| **edge recall** | 0.970 | 0.996 |
| **edge f-1 score** | 0.969 | 0.966 |

Supplementary Table 2: Evaluation of the LSTree VS Elephant lineage prediction for mouse embryo data.

### Supplementary Note 6: Fixation and backtracking challenges

At any moment during live imaging it is possible to stop the recording and perform fixation and immunofluorescence steps in order to get more functional information on the developmental state of the cells. Considering that all necessary solutions are prepared ahead of time, the added amount of time for having a light-sheet recording + fixation step is usually of maximum 24 hours (as this clearly depends on the time used for each immunofluorescence step). The main challenge associated with fixation of samples in Matrigel is the resulting structural change of the initial dome, which most often causes the organoids to sink to the bottom of the holder. Therefore we have performed these steps typically in the microscope in order to keep track of the sample of interest as the drop collapses. This is a very fast procedure, and we noticed that typically most major movements occur between minutes 5 and 10 after fixative addition. Therefore, to keep track of possible organoid drifts, we change the stack dimensions during fixation to accommodate a larger imaging volume (exact dimensions vary but could be up to 40 planes every 10 μm). After staining is done, we revert to original stack dimensions to aid the registration step. An example of the registration parameters is provided in ‘Elastix_parameter_Affine.txt’ file in the Source Code.

To avoid unnecessary computational times, registration is done by mapping the fixed stack onto the last timepoint of the live recording. Since the live recording needs to be stopped in order to start fixation, the most crucial timing involves the fixation of the sample itself, so that we can still keep track of shape and cellular organization. Using 4% PFA we noticed samples becoming fixed 10 minutes after adding the fixative, meaning that the timepoint after fixation is usually happening at a slightly longer temporal spacing than that during the live recording. Still, for spherical samples this is still good enough to register with a good approximation the positions of the cells before and after fixation. To mention, not all organoids will require a registration step to bridge live imaging with fixation volumes, as this will highly depend on the position of the organoid in the drop. For the future. new fixation protocols may aid in order to avoid Matrigel drop collapse. This will prevent organoids to change their relative position during fixation, thus minimizing the requirement of dedicated registration steps, and improving overall experiment efficiency. We have noted that the addition of Glutaraldehyde to the PFA might help in maintaining organoids in place. Nonetheless, we envision that, apart from using different fixative for this step, one of the main challenges may be that the fixation protocols still allow proper staining with off-the-shelf antibodies.

### Supplementary Tables

**Supplementary Table 1: List of extracted features Figure 1f**

| **Feature name** | **Short description** |
| --- | --- |
| Nuclei number | number of nuclei inside epithelium |
| Epithelium volume | (organoid – lumen) segmentation volumes |
| Nuclei density | nuclei number / epithelium volume |
| Mean cell volume | cell volume / cell number |
| Mean cell / nuclei volume | mean cell volume / mean nuclei volume |

**Supplementary Table 2: List of extracted features**

| **Feature name** | **Short description** |
| --- | --- |
| gen#_label | label of all progeny of #-generation |
| time_since_split | time elapsed since last division |
| displacement | spatial displacement between two consecutive nodes |
| nuclei_intensity_* | * evaluation of nuclei intensity |
| nuclei_**_radius | ** radius of the segmented nucleous |
| nuclei_volume | volume of the segmented nucleous |
| nuclei_dist_to_***_mean | men distance of nucleous pixels to *** boundaries |
| cell_**_radius | ** radius of the segmented cell |
| cell_volume | volume of the segmented cell |
| cell_neighbors | estimated first neighbors of particular cell |
| merge | point where two tracklets fuse into one |
| merged_branch_id | node ID’s from sister branches that end in a merge |
| merged_track | nod ID’s of all subsequent tracks directly linked to a merge |

* can be *mean*, *std* (standard deviation), and quartile-based ranging from *0.00 , 0.25, 0.50, 0.75* and *1.00*

** *mean, maximum or minimum*

*** *basal, lumen*

### Supplementary Figures

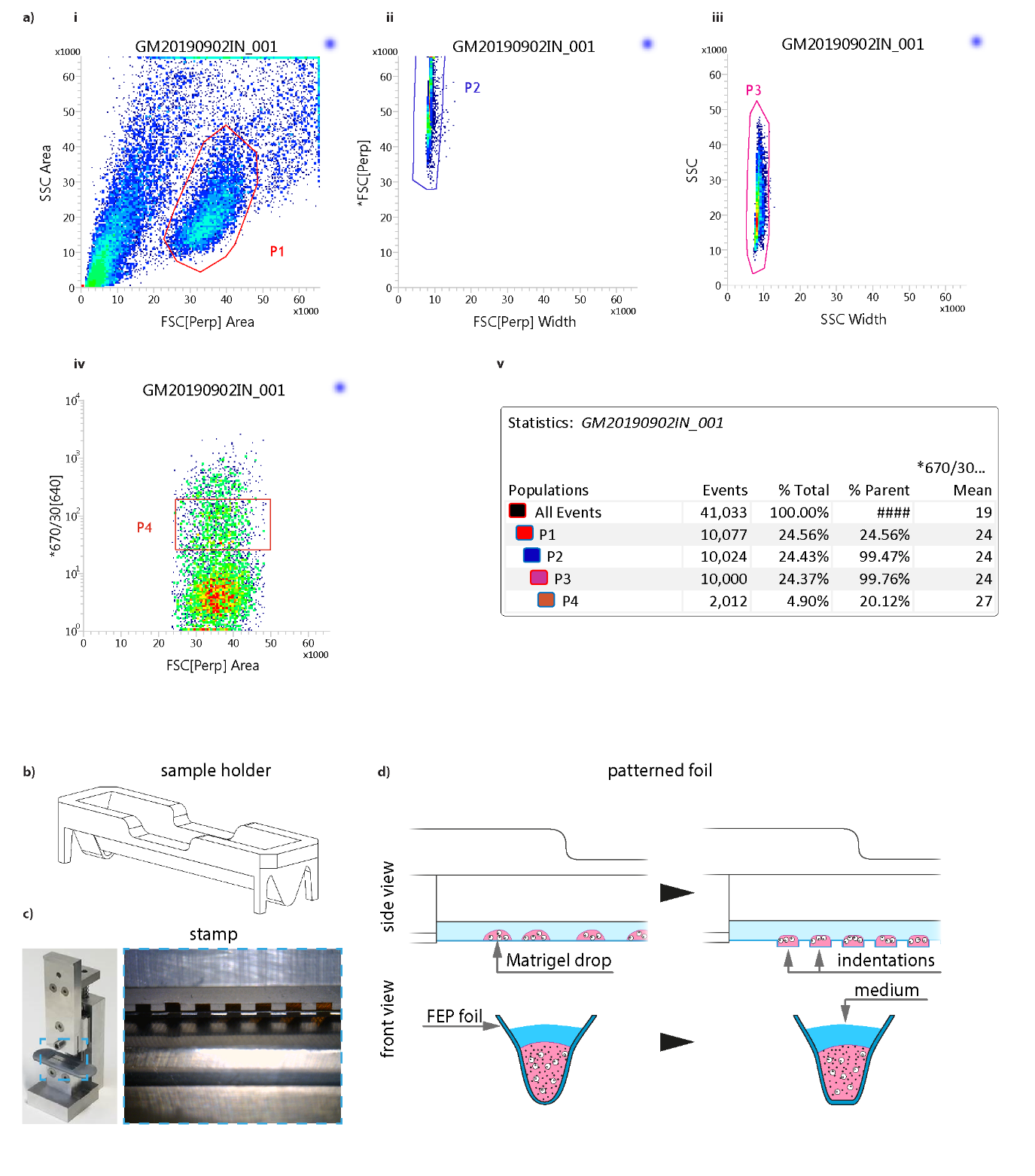

**Supplementary Figure 1**

**Gating strategy and sample holder geometry.**

**a)** Gating strategy for all FACS sorting. i) Sorting based on forward vs side scattering to take out debris, ii,iii) second and third gates ensure the filtering of doublets from the sample taking into account forward and side scatter and event width, iv) fluorescence intensity gating is done in such a way to avoid the highest expressing cells, as these can have multiple insertions of the construct and thus not develop properly, and v) statistics of the gating strategy. At the end of the sort, 83700 miRFP+ cells were found. **b)** Perspective view of the sample holder. **c)** Pictures of the aluminum stamp used for creating the compartments onto the FEP foil. **d)** Side and front views of part of the sample holder, depicting the FEP foil (dark blue line), the Matrigel drops (pink) with cells and the medium (blue), without (left) or with (right) stamp patterning. The dimension of each compartment is 1.5X1.5 mm^2^ and 1 mm deep.

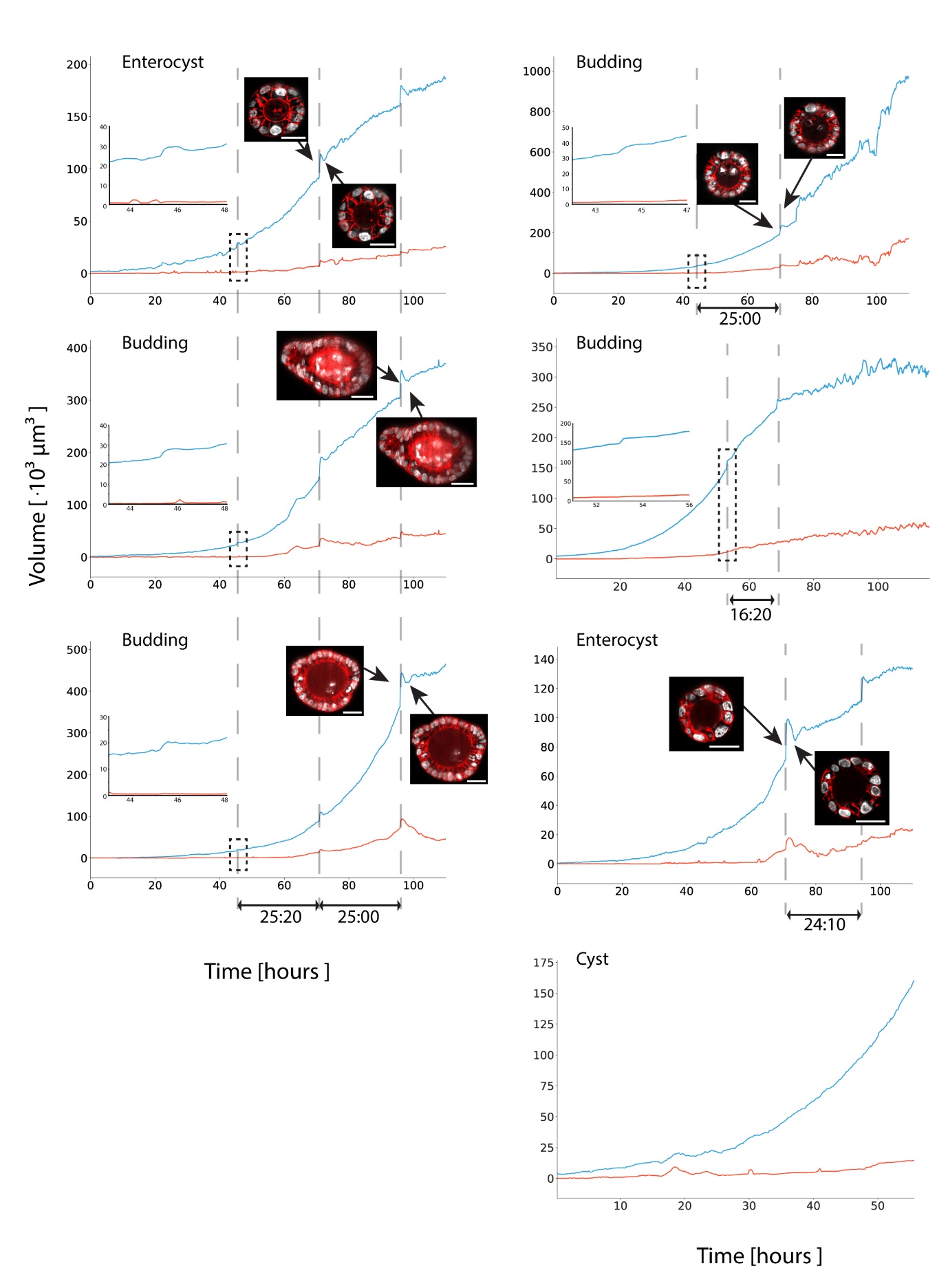

**Supplementary Figure 2**

**Organoid and lumen volume dynamics.**

Organoid and lumen volumes for all datasets. Sudden volume increase is related to moments of medium exchange, having a direct impact on an increase in image quality, which are also recognized in the H2B-mCherry (gray) / mem9-GFP (red) data snippets. All scale bars depict 25 μm.

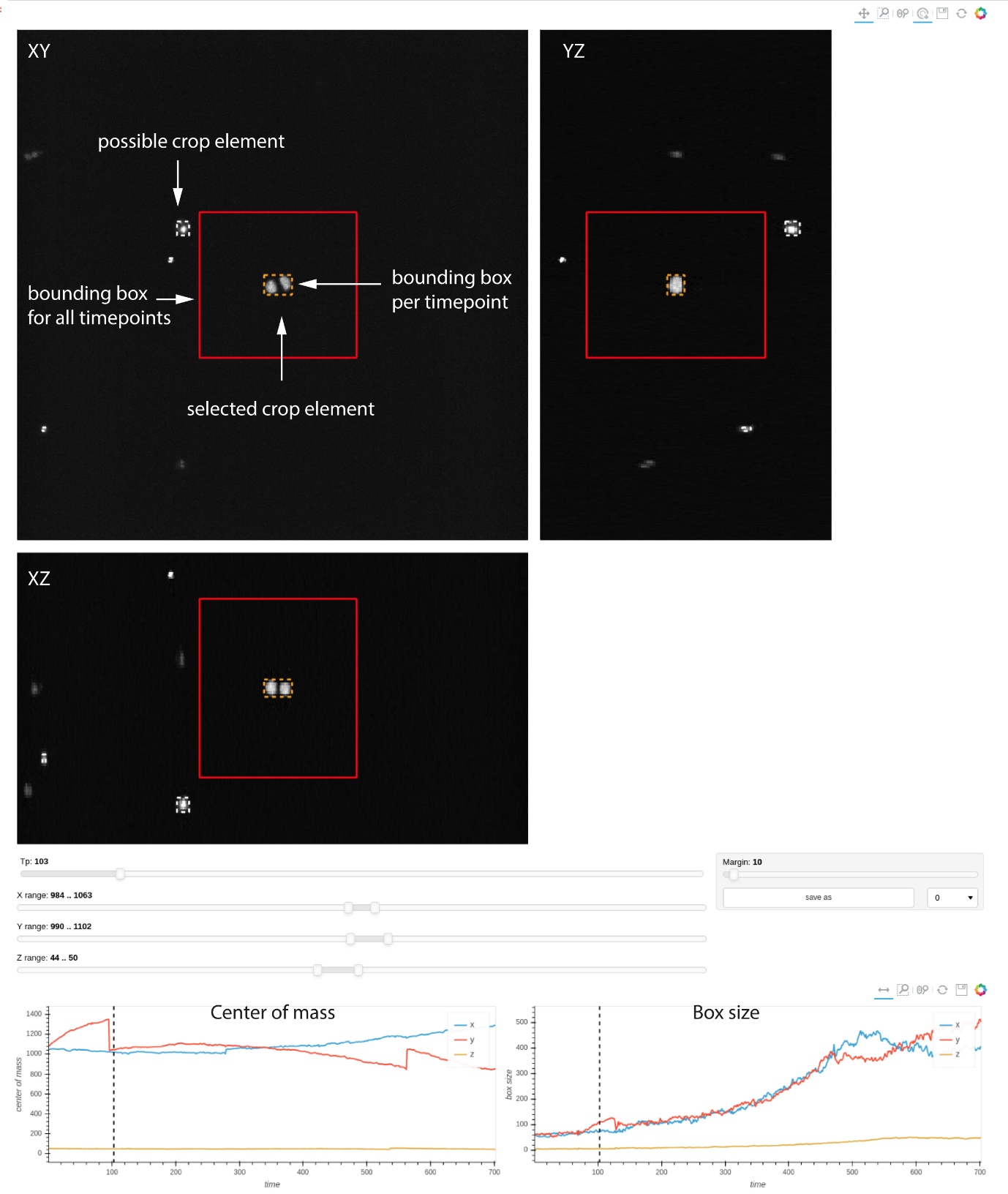

**Supplementary Figure 3**

**Cropping and registration pre-processing.**

Example screenshot of the crop correction tool in LSTree. Top: XY, XZ and YZ views of a certain timepoint of the raw data. All possible crop elements (organoids) are shown with a dashed box surrounding them, and only the selected element shows the respective current timepoint bounding box (orange), and the maximum bounding box (red). Bottom: Center of mass (left) and box size (right) plots for each spatial dimension throughout the entire movie. Sudden jumps in values for each of these graphs can point to either correct reassignments of the bounding boxes due to e.g. medium change leading to organoids jumps around the FOV, as well as to possible moments where correction might be needed.

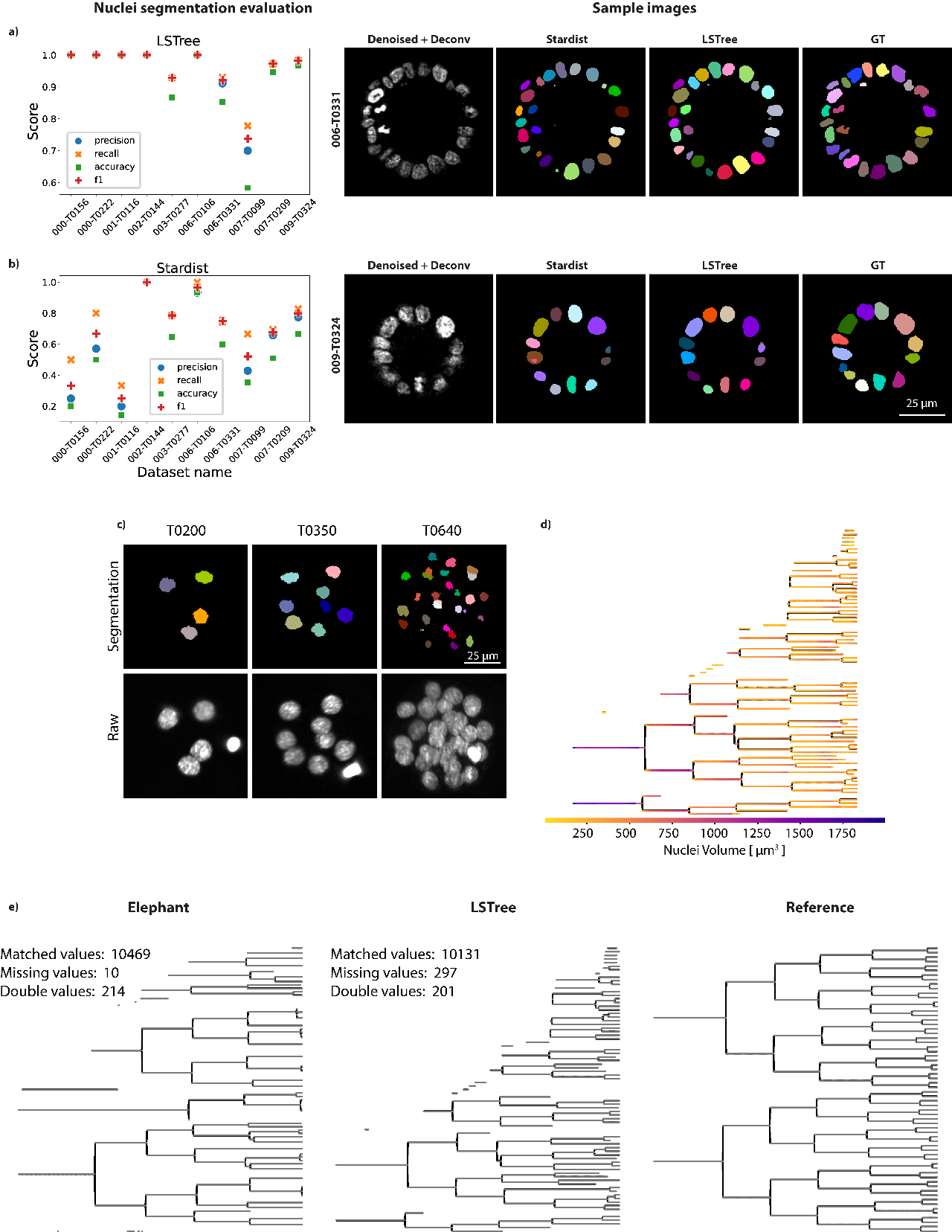

**Supplementary Figure 4**

**Assessing quality of nuclei segmentation and tree prediction strategies.**

**a,b, left)** Evaluation of nuclei segmentation with our modified version of the RDCNet and of Stardist **right)** Example images of two stacks used for evaluation. GT, grount-truth**. c)** nuclei segmentation prediction using background subtracted raw data from previous publication. Worse segmentation at later timepoints related to worse image quality. **d)** Tree prediction showing nuclei volume values based on segmented nuclei. **e)** Validation and comparison of lineage tree prediction using Elephant (left) and LSTree (center) in respect to ground truth from previous publication (right).

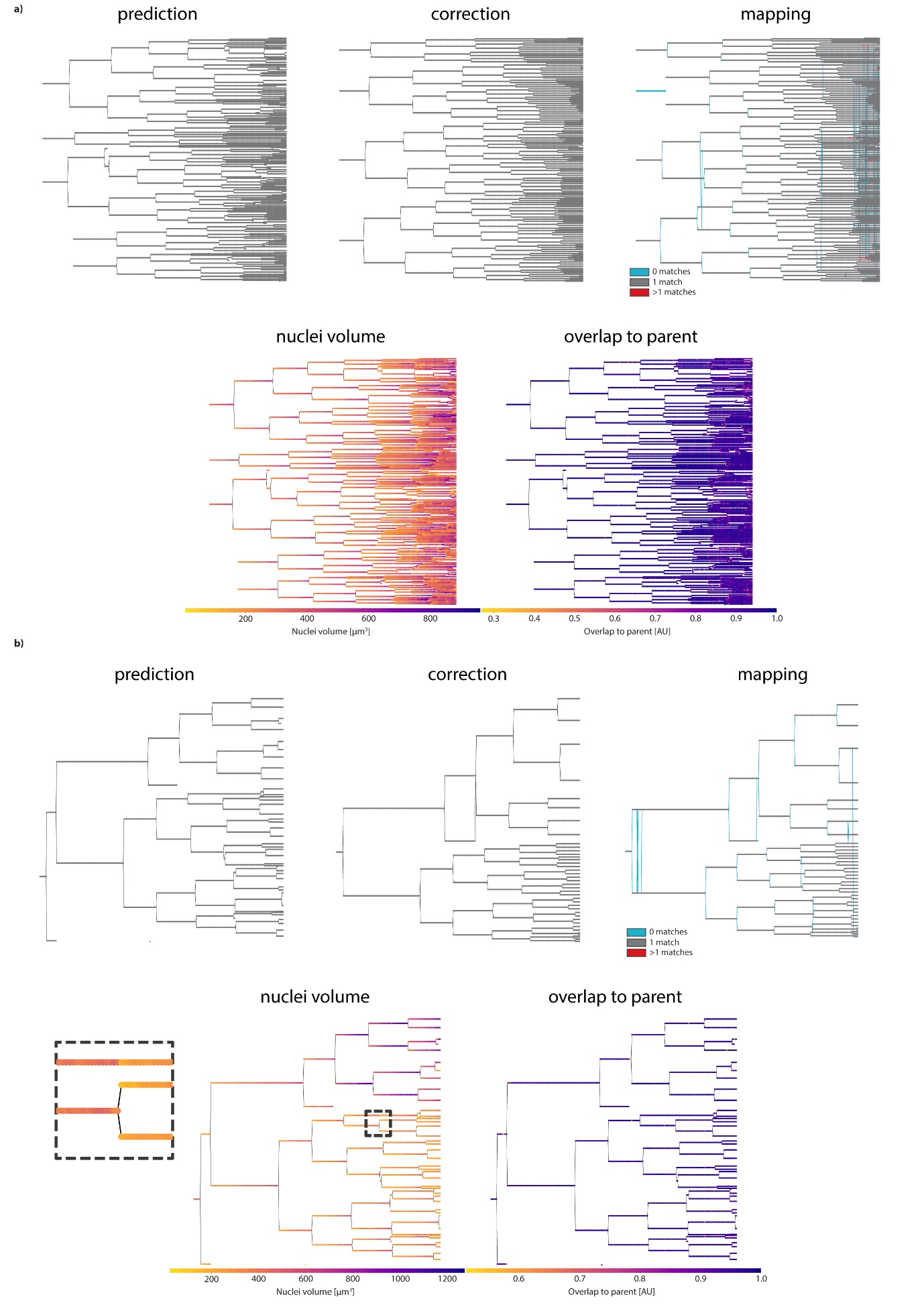

**Supplementary Figure 5**

**Lineage tree prediction evaluation methods.**

**a)** Above: prediction, correction and mapping trees for one of the budding datasets used for validation of the tree prediction network. Mapping tree is the predicted tree mapped onto the correct one, with 0 matches meaning missing nodes (related to missing spots), 1 match corresponds to correct prediction and 2 or more matches relate to nodes where 2 or more spots have been assigned to the same node. Below: Nuclei volume and overlap to parent features overlaid on the predicted tree to aid curation. Overlap to parent typically becomes less robust as the organoid grows, since cell packing and lower image quality hinders a better overlap assignment. **b)** Same as in a) for the second validation dataset. Here it is interesting to note the missing daughter in highlighted in the dashed box. Nuclei volume are particularly helpful when recognizing missing divisions, as a sudden volume change on the same track is readily seen.

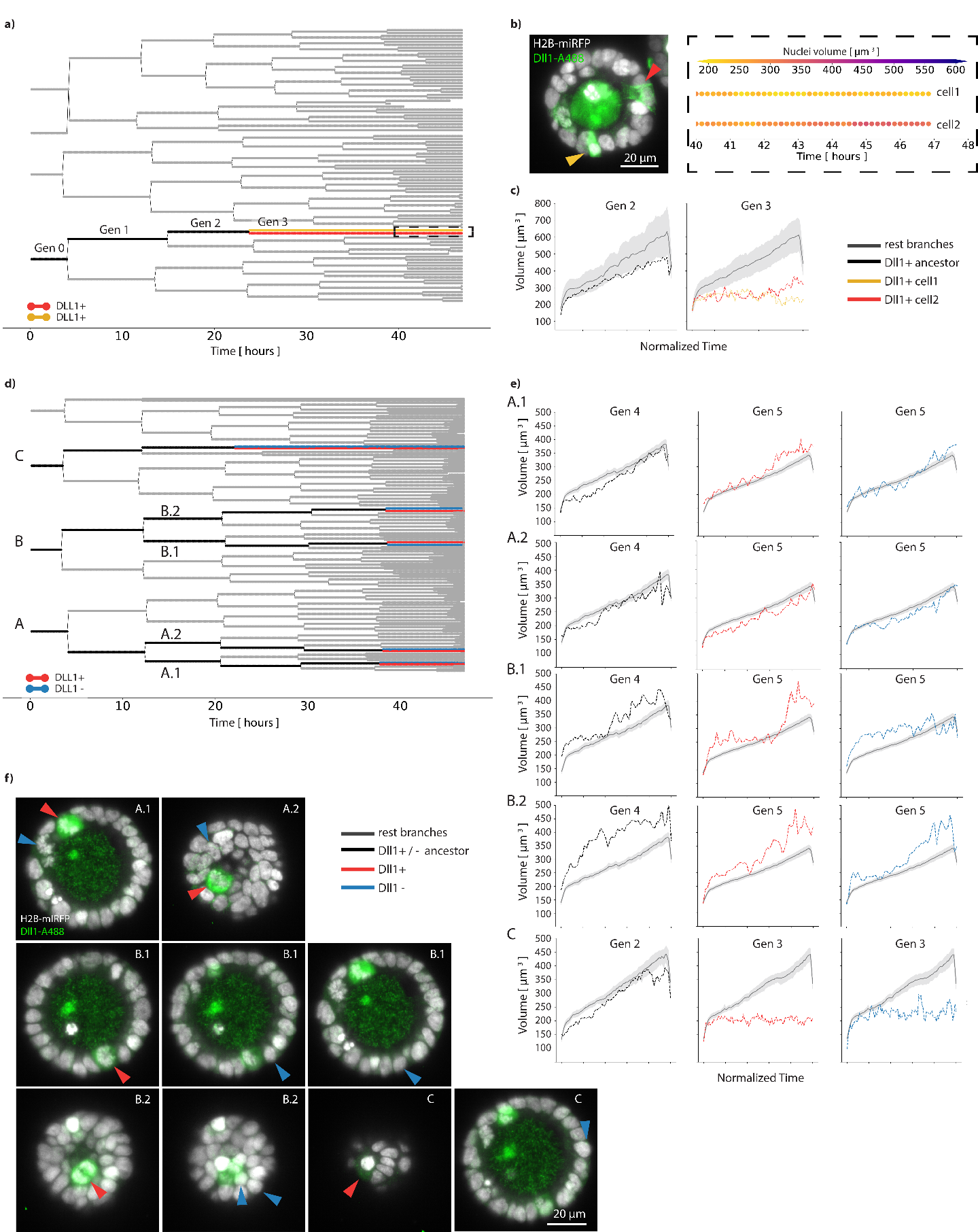

**Supplementary Figure 6**

**Backtracking after fixation and staining.**

**a)** Backtracking tree where two sister cells stop dividing at generation 3, both being DLL1+ at the end of the recording. **b) Left:** Cross-section image of cyst after fixation and staining with DLL1 antibody. Arrow colors match backtracking colors in a). **Right:** Expanded dashed selection box from a) showing both sister cell tracks overlaid with their nuclei volume information. Cell2 has larger nuclei volume than cell1 at the end of the recording. **c)** Nuclei volume distribution per generation. It is important to note that the tendentially larger nuclei volume from the rest branches follows 1) the trend that during generation 3 all branches apart from the backtracked ones went over full cell cycle and 2) that this lineage tree has a region dominated by an early binucleation event, leading to larger nuclei volume over all generations. **d)** Second backtracking example, showing multiple cells backtracked. DLL1+, DLL1- cells shown in red and blue respectively. Ancestors in black. **e)** Nuclei volume plots for each end generation of ancestor, DLL1+ and DLL1- branches. Color coding following backtracking scheme in d). **f)** Cross sections depicting the stained cells for DLL1 at the end of the live recording. For all plots, whenever more than one track is being evaluated for the same label, the full line represents the mean whereas the shaded region corresponds to 95% of the confidence interval. A total of 4 different experiments were performed with similar results. Source data are provided as a Source Data file.

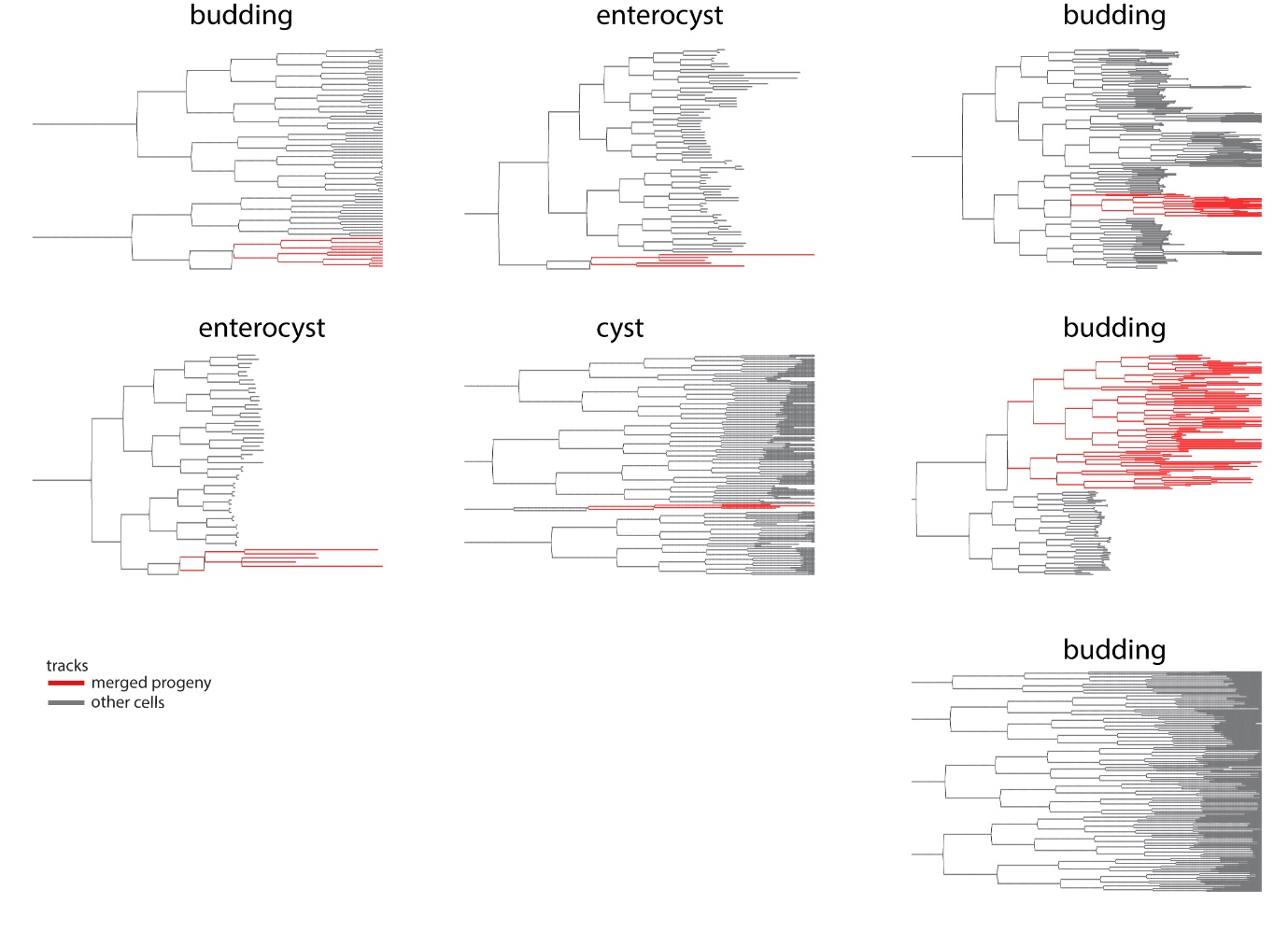

**Supplementary Figure 7**

**Binucleation events during organoid growth.**

Summary of all lineage trees used in this work, highlighting the progeny of the early mitotic division failure in red against all other tracks (grey). Note the lower right “Budding” tree, which is the only dataset not showing merging events.

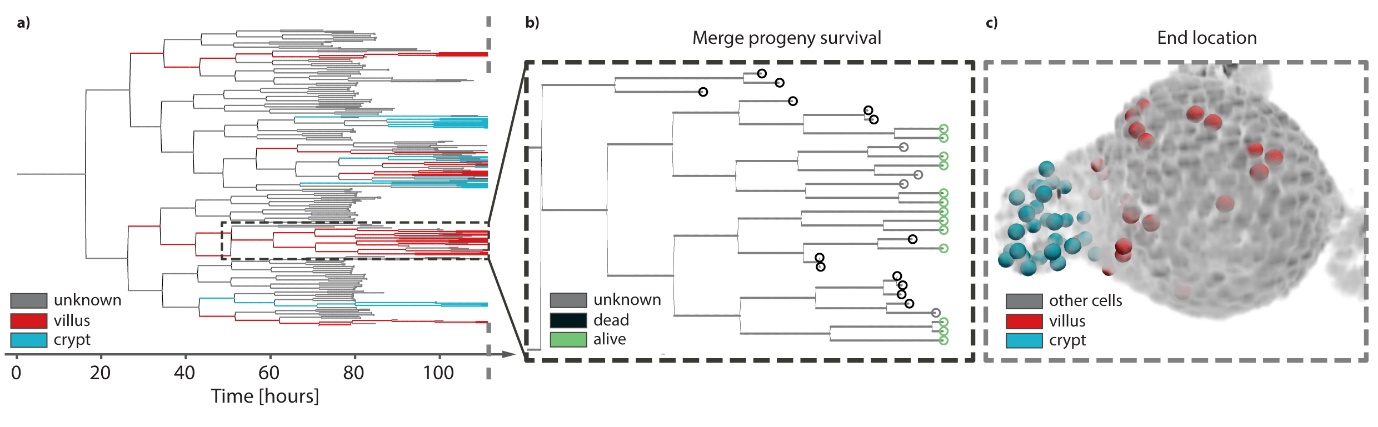

**Supplementary Figure 8**

**Localization of binucleated progeny.**

**a)** Lineage tree of budding organoid depicting cells that end up in the villus (red), in the crypt (cyan) or have unknown end location (grey). Notice that all cells in the dashed box correspond to the merge progeny and all end up in the villus region, whereas cells from other tree regions may form the crypt. **b)** Detail of the dashed box in a), depicting the tracks that end with dead (black), unknown (grey) or alive (green) cells. Dead cells are typically extruded into the lumen, unknown cells represent nuclei that were not trackable anymore due to e.g. poor image quality and high cell packing, and alive cells are nuclei that remain part of the epithelium until end of the recording. **c)** 3D representation of raw (nuclei) data (gray) overlayed with the cells from the tracks presented in a) at the last timepoint, depicting the distribution of crypt and villus cells found around the organoid.

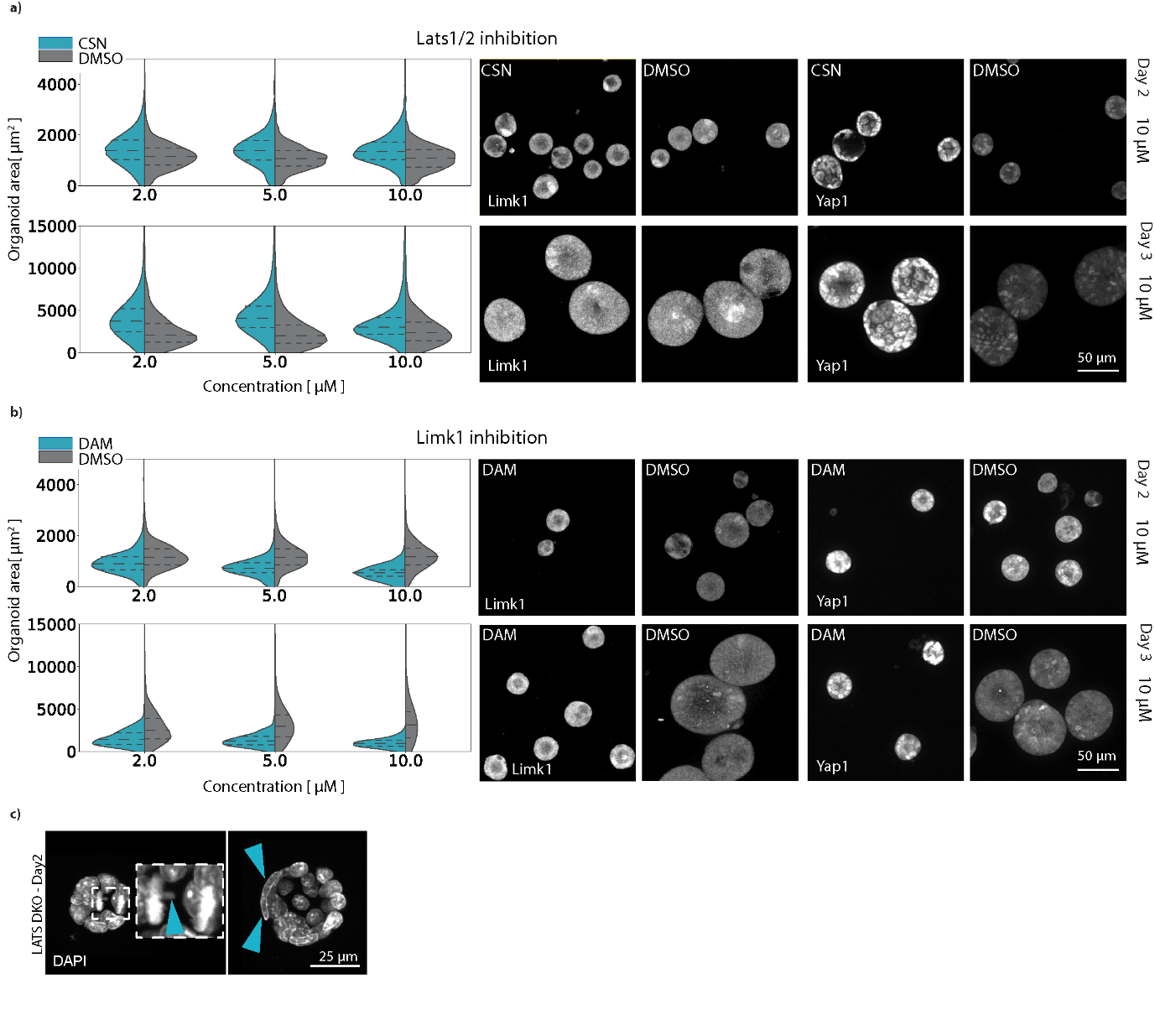

**Supplementary Figure 9**

**Limk1 or Lats1/2 inhibition influence organoid growth.**

**a)** LEFT: Organoid volume distribution for different concentrations of the Lats1/2 inhibitor used against DMSO control ($n_{CSN, day2}=4161$, $n_{CSN, day3}=5389$, $n_{DMSO, day2}=4279$ and $n_{DMSO, day3}=3277$). RIGHT: representative images of Limk1 and Yap1 signal for both control and compound (at 10μM concentration) conditions. **b)** LEFT: Organoid volume distribution for different concentrations of the Limk1 inhibitor used against DMSO control ($n_{DAM, day2}=2528$, $n_{DAM, day3}=3868$, $n_{DMSO, day2}=3713$ and $n_{DMSO, day3}=3157$). RIGHT: representative images of Limk1 and Yap1 signal for both control and compound (at 10μM concentration) conditions. Shown is data for Days 2 and 3 of fixation. **c)** Representative images of cysts at Day2 from a LatsDKO organoid line. Mitotic errors and binucleated cells highlighted with blue arrows. The dashed lines inside each plot correspond to the first and third quartile of the values from either CSN or DAM treated organoids, with the median as the dashed line in between them. Experiment was performed with a replicate, showing similar results. Source data are provided as a Source Data file.
